# Functional synaptic connectivity of engrafted spinal cord neurons with locomotor circuitry in the injured spinal cord

**DOI:** 10.1101/2025.04.05.644402

**Authors:** Ashley Tucker, Angelina Baltazar, Jaclyn T. Eisdorfer, Joshua K. Thackray, Katie Vo, Hannah Thomas, Avnika Tandon, Joshua Moses, Brendan Singletary, Tucker Gillespie, Ashley Smith, Anna Pauken, Sneha Nadella, Michael Pitonak, Sunjay Letchuman, Julius Jang, Michael Totty, Frank L. Jalufka, Miriam Aceves, Andrew F. Adler, Stephen Maren, Heath Blackmon, Dylan A. McCreedy, Victoria Abraira, Jennifer N. Dulin

## Abstract

Spinal cord injury (SCI) results in significant neurological deficits, with no currently available curative therapies. Neural progenitor cell (NPC) transplantation has emerged as a promising approach for neural repair, as graft-derived neurons (GDNs) can integrate into the host spinal cord and support axon regeneration. However, the mechanisms underlying functional recovery remain poorly understood. In this study, we investigate the synaptic integration of NPC-derived neurons into locomotor circuits, the projection patterns of distinct neuronal subtypes, and their potential to modulate motor circuit activity. Using transsynaptic tracing in a mouse thoracic contusion SCI model, we found that NPC-derived neurons form synaptic connections with host locomotor circuits, albeit at low frequencies. Furthermore, we mapped the axon projections of V0C and V2a interneurons, revealing distinct termination patterns within host spinal cord laminae. To assess functional integration, we employed chemogenetic activation of GDNs, which induced muscle activity in a subset of transplanted animals. However, NPC transplantation alone did not significantly improve locomotor recovery, highlighting a key challenge in the field. Our findings suggest that while GDNs can integrate into host circuits and modulate motor activity, synaptic connectivity remains a limiting factor in functional recovery. Future studies should focus on enhancing graft-host connectivity and optimizing transplantation strategies to maximize therapeutic benefits for SCI.

## Introduction

Spinal cord injury (SCI) causes profound neurological deficits that vary in severity based on the level and extent of the injury including paralysis, dysesthesia, pain, and autonomic dysfunction. An estimated 300,000 individuals in the United States are living with SCI, with approximately 17,000 new cases occurring each year, most often resulting from motor vehicle accidents and falls^1^. Treatment is focused on improving quality of life through rehabilitation and lifestyle modification, but no curative therapy is available to date. Through decades of preclinical research, we now appreciate that engrafted NPCs differentiate into glial cells and diverse subtypes of neurons^2–9^, and that grafts support host axon regeneration and establish new synaptic connections with the injured host nervous system^5,8,10–17^. Some studies have reported recovery of locomotor function following NPC transplantation in rodent SCI models^5,18–23^. Notably, NPC transplantation has been evaluated in multiple human clinical studies including an ongoing trial to evaluate human induced pluripotent stem cell (hiPSC)-derived NPCs for subacute SCI^24,25^. Together, these observations highlight NPC transplantation as a promising therapeutic approach to improve locomotor function following SCI.

Despite this promise, there remain knowledge gaps that act as barriers to the clinical efficacy of NPC transplantation. Functional recovery in transplanted animals is typically modest, and reports of locomotor recovery have been challenged at times by failed reproducibility studies^26,27^. Importantly, there is little mechanistic understanding of how engrafted neurons can impart functional benefits. Although it is known that transplant-derived neurons can synaptically integrate with the host nervous system, it is unclear if this synaptic integration occurs in a biologically relevant manner. It is also unclear whether functional recovery is a result of activity through graft-mediated neural relays, and/or via another “paracrine” function of graft neurons in parallel to this connectivity. It is unknown whether specific phenotypes of transplant-derived neurons can impart greater functional benefits than others. Finally, it is unclear whether graft-host connections need to be stabilized and strengthened via biologically relevant “training”; e.g., through activity-based rehabilitation. Addressing these critical knowledge gaps can accelerate the realization of the therapeutic potential of neural replacement strategies in SCI and in other indications.

Our understanding of the mechanisms through which NPCs synaptically and functionally integrate with multiple neural pathways to restore motor function remains limited. In this study, we investigate how NPCs synaptically integrate into locomotor circuitry, how different subpopulations of NPCs project within the host spinal cord and form synapses, and how the activity of distinct graft cell subtypes influences motor circuit activity.

## Results

### Multiple phenotypes of NPC graft-derived neurons can synaptically integrate with spinal locomotor circuitry

Several studies have reported improvements in locomotor function following NPC transplantation into the injured spinal cord^5,19–23^. It is generally thought that graft-derived neurons (GDNs) support motor functional recovery via formation of new graft/host neural relay*s*^28,29^. However, the extent to which GDNs can spontaneously synaptically integrate into spinal locomotor circuitry has not yet been investigated. To address this, we performed transsynaptic tracing experiments to map the synaptic connections between GDNs and host spinal locomotor circuits in mice. We utilized two different transsynaptic tracing strategies in this study: monosynaptic tracing with the G-deleted, EnvA rabies virus (RABV)^30–32^, and polysynaptic circuit mapping using the Bartha strain of pseudorabies virus^33,34^. The NPCs used in our study are derived from E12.5 mouse spinal cords; these primary rodent cells are useful for addressing the fundamental biology of graft/host interactions because they are developmentally programmed to differentiate into multiple subtypes of spinal cord neurons^7,35,36^ and the use of inbred mouse strains allows for syngeneic grafting studies.

We first performed monosynaptic tracing to characterize the numbers and types of synaptic connections that are spontaneously established between GDNs and host lumbar motor neurons. We initiated Cre-dependent RABV tracing from spinal lumbar motor neurons in adult ChAT-cre mice at 8 weeks following transplantation of GFP^+^ NPCs into sites of contusion SCI at spinal level T12, then sacrificed for histological analysis two weeks later (**Fig. 1a**). We selected ChAT-cre mice as a driver for motor neuron-specific gene expression based on our observations that Hb9-cre mice exhibited off-target gene expression in cells other than spinal motor neurons^37^ (**Fig. S1**). We verified that RABV^+^ expression was restricted to ChAT^+^ motor neurons in the ventral spinal cord gray matter of uninjured adult ChAT-cre mice (**Fig. 1b-c**). At ten weeks following NPC transplantation into sites of SCI in ChAT-cre mice, RABV^+^ neurons were distributed throughout grafts of some, but not all subjects (**Fig. 1d**). Subsets of these RABV^+^ cells expressed markers of excitatory neurons (CamKII, **Fig. 1e**), cholinergic neurons (ChAT, **Fig. 1f**), and GABAergic neurons (GAD65/67, **Fig. 1g**). Quantification revealed that there were no significant differences in the abundances of each phenotype of RABV^+^ neuron in grafts. Moreover, 9 out of 22 animals assessed did not have any RABV^+^ neurons in grafts (**Fig. 1h**). These data suggest that the spontaneous degree of synaptogenesis between GDNs and host locomotor circuitry is very low, with an average of only 0.018 ± 0.0057% of graft neurons labeled with rabies. Because the size and shape of each graft is variable, we asked whether the abundance of RABV^+^ neurons in grafts is correlated to graft volume; we found no significant correlation (**Fig. 1i**).

**Figure 1.**
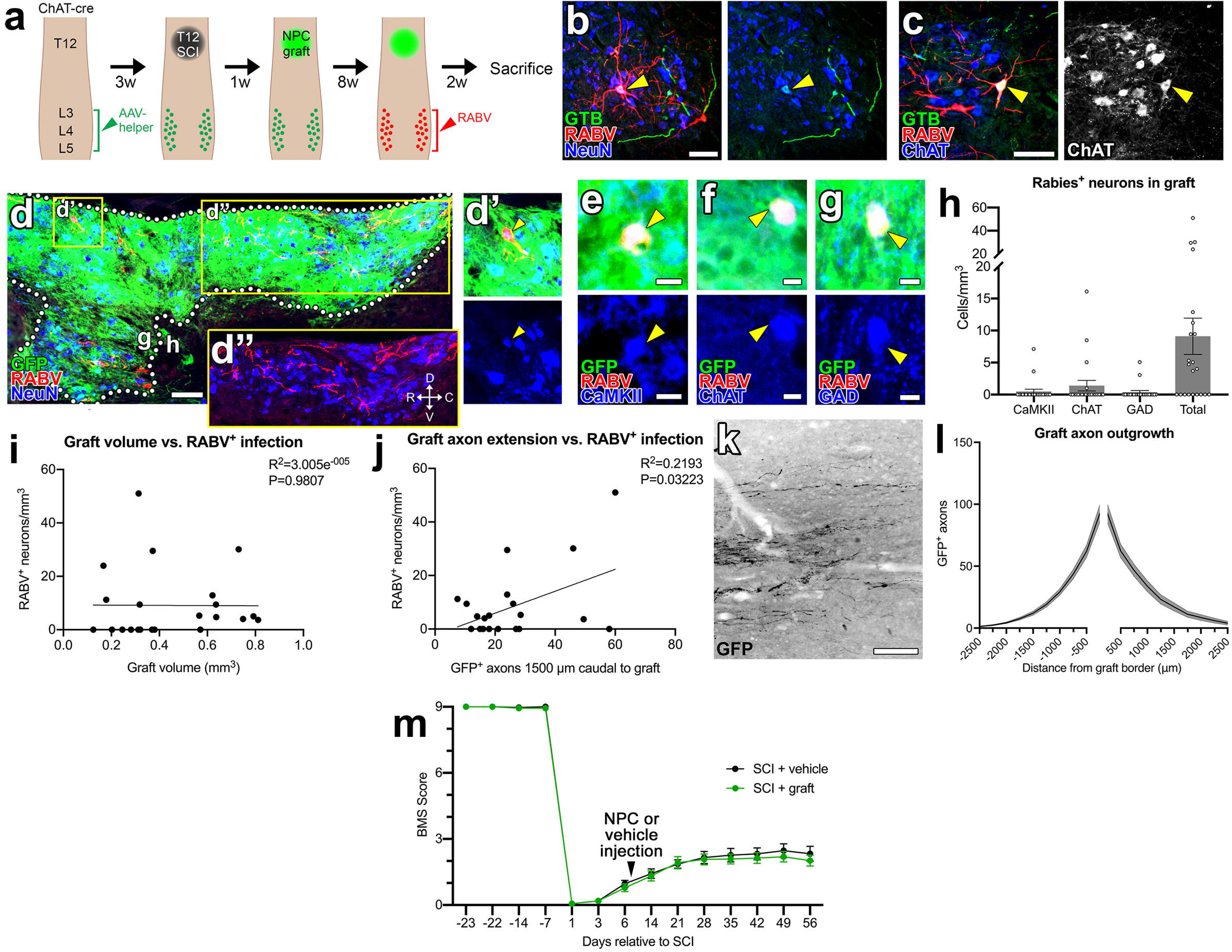
Monosynaptic rabies tracing reveals a small percentage of graft-derived neurons with monosynaptic connections onto lumbar spinal cord motor neurons. (**a**) Experimental strategy for monosynaptic rabies tracing. AAV2 encoding Cre-dependent rabies helper components was injected into ventral horns of the L3-L5 spinal cord of adult ChAT-cre mice. Three weeks later, a contusion injury was administered at spinal level T12; 1 week after injury, GFP^+^ NPCs were transplanted at the lesion site; 8 weeks after transplantation, G-deleted, EnvA SAD-b19 rabies-mCherry (RABV) was injected into the L3-L5 spinal cord, and mice were sacrificed 2 weeks later. The same subjects also underwent open field locomotor (Basso Mouse Scale) testing throughout the duration of the study, until immediately prior to virus injection. (**b, c**) Images of RABV expression in the L4 spinal cord of uninjured ChAT-cre mice. Arrowheads show colocalization of RABV with rabies helper components (GTB) in (**b**) NeuN^+^ neuron and (**c**) ChAT ^+^ motor neuron. (**d**) Image of GFP^+^ NPC graft with RABV^+^ neurons (red). The graft (g) /host (h) boundary is shown with dotted lines. Arrowhead in **d’** indicates one example of a RABV^+^/NeuN^+^ graft cell. Panel **d”** shows graft without GFP immunofluorescence. Rostral (R), caudal (C), dorsal (D), and ventral (V) directions are indicated with arrows. (**e-g**) High-magnification images of RABV^+^ graft neurons expressing (**e**) CaMKII, (**f**) ChAT, or (**g**) GAD65/67 (arrowheads). (**h**) Quantification of the density of graft RABV^+^ neurons that express CaMKII, ChAT, or GAD65/67; the total density of RABV^+^ graft neurons per subject is shown for n=22 subjects. (**i**) Correlation of graft volume vs. the density of RABV^+^ neurons in the graft for n=22 subjects. (**j**) Correlation of the total number of graft axons extending to 1500 μm caudal to the graft/host border vs. the density of RABV^+^ neurons in the graft for n=21 subjects (one subject was excluded from axon quantification analysis due to poor tissue quality). (**k**) Example image of graft axon outgrowth at 1 mm caudal to the graft/host border. (**l**) Quantification of total graft axon outgrowth at 250-μm intervals rostral and caudal to the graft/host border for n=29 subjects. (**m**) Basso Mouse Scale scores for 41 mice (n=16 SCI + vehicle; n=25 SCI + NPC graft) up to 8 weeks post-injury. All data are mean ± SEM. Scale bars = 100 μm (**b, c, d, k**), 10 μm (**e, f, g**).

However, we did detect a significant positive correlation between graft RABV^+^ neuron density and graft axon extension 1.5 mm into the caudal spinal cord (**Fig. 1j**). It is not surprising that the degree of graft connectivity is directly correlated with the ability of the graft to extend axons throughout the host spinal cord. However, the degree of spontaneous axon extension from NPC grafts is modest, with only a few hundred axons extending per graft (**Fig. 1k-l**). This is consistent with our previous findings in mouse NPC grafting models^9,35,38^.

Previous work has shown that NPC transplantation can promote locomotor recovery after SCI^5,18–23^. However, not all studies have reported positive effects of grafting on locomotor recovery^26,27,35^. To determine whether NPC transplantation can improve locomotor recovery in our SCI model, we conducted open field locomotor assessments in the same animals, for 8 weeks post-transplantation (prior to virus injection). Strikingly, we failed to observe significant differences in locomotor function between grafted and non-grafted animals (**Fig. 1m**; n=16 SCI + vehicle; n=25 SCI + NPC graft; p=0.992).

Considering the possibility that GDN synaptic integration with host locomotor circuits might occur via polysynaptic relays, we took a parallel approach to neural circuit mapping using pseudorabies virus (PRV). PRV tracing has previously been utilized to visualize plasticity after SCI and to map graft/host connectivity following NPC transplantation into the injured spinal cord^10,39–43^. We injected PRV-RFP directly into the sciatic nerves of injured and transplanted mice, then assessed histological outcomes 72 h later (**Fig. 2a**). PRV-RFP expression was evident in lumbar motor neurons and spinal cord interneurons at 72 h post-injection (**Fig. 2b; Supp. Movies 1-2**), validating that intrasciatic delivery was sufficient to transduce MNs. In injured/grafted subjects, PRV was detectable in neurons throughout the host spinal cord rostral and caudal to the lesion site as well as within the graft itself (**Fig. 2c**). PRV spreads polysynaptically, so GDNs expressing PRV-RFP are either directly or indirectly integrated into locomotor circuits. We performed immunohistochemistry to characterize the phenotypes of GDNs that were labeled with PRV and found that all three major neurotransmitter phenotypes (**Fig. 2d-f**), calbindin^+^ and parvalbumin^+^ V1 interneurons (**Fig. 2g-h**), and Chx10^+^ V2a interneurons (**Fig. 2f**) were represented. Notably, we observed 20-fold more PRV-labeled GDNs compared to RABV-labeled GDNs (0.43% PRV^+^ vs. 0.02% RABV^+^). This indicates that GDNs are more likely to integrate into polysynaptic relay circuits with spinal locomotor circuitry, rather than making direct synaptic connections. Taken together, these data reveal that the percentage of GDNs that spontaneously integrate into spinal locomotor circuits are quite low, representing less than 1% of all graft neurons.

**Figure 2.**
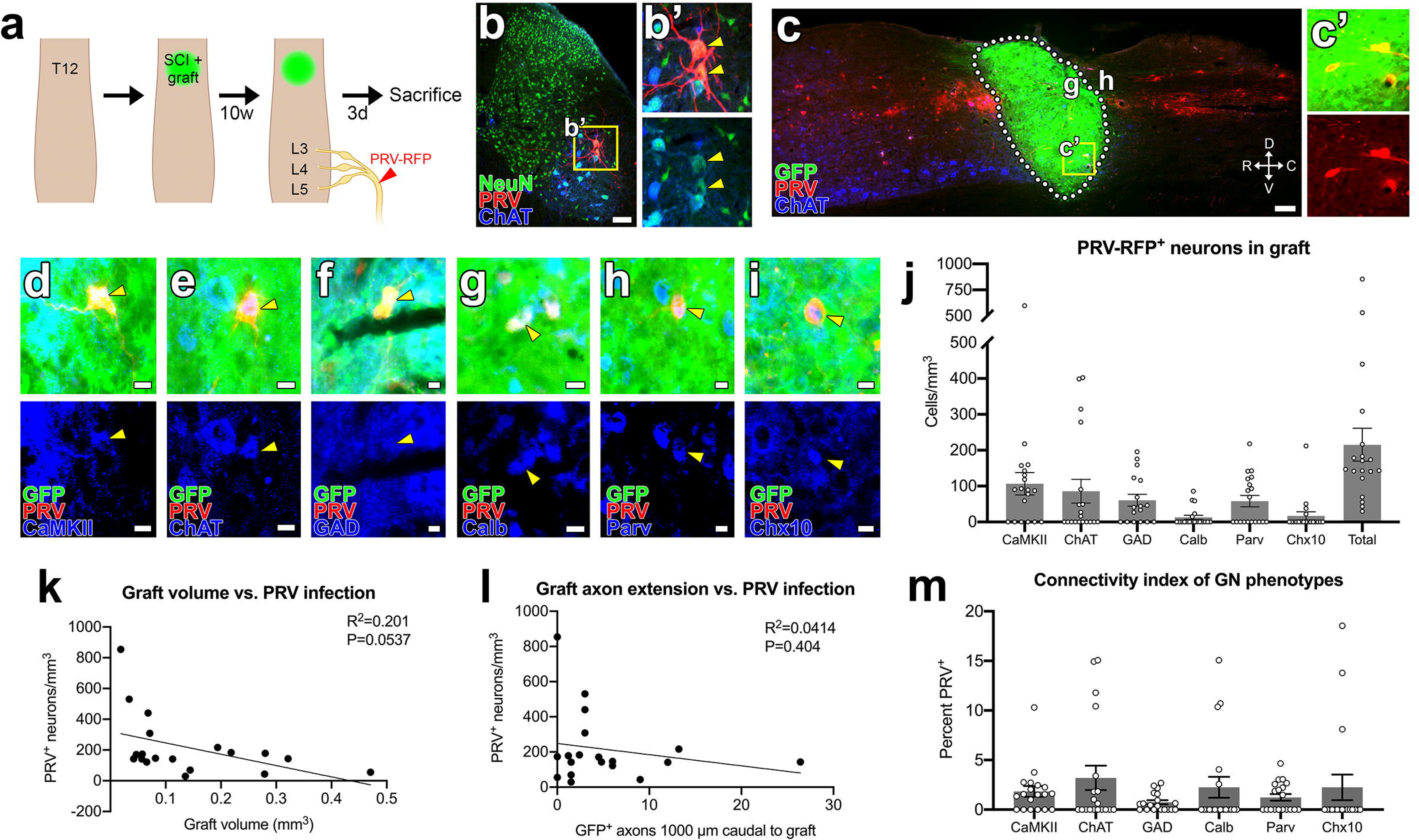
Polysynaptic tracing from the sciatic nerve labels multiple subtypes of graft-derived neurons. (**a**) Experimental strategy for pseudorabies tracing. Dorsal column lesion SCI was administered at spinal level T12, followed by immediate transplantation of GFP^+^ NPCs into the lesion cavity. Ten weeks later, PRV-RFP was injected into bilateral sciatic nerves; three days after PRV injection, animals were sacrificed for histological analysis. (**b**) Transverse image of the L4 spinal cord at 60 hours after PRV-RFP injection into the sciatic nerve. (**b’**) High-magnification image of PRV^+^/ChAT^+^ motor neurons in the ventral horn (arrowheads). (**c**) Sagittal image of a GFP^+^ NPC graft containing PRV^+^ neurons (**c’**). The graft (g) /host (h) boundary is shown with dotted lines. Rostral (R), caudal (C), dorsal (D), and ventral (V) directions are indicated with arrows. (**d-i**) High-magnification images of PRV^+^ graft neurons expressing (**d**) CaMKII, (**e**) ChAT, (**f**) GAD65/67, (**g**) calbindin, (**h**) parvalbumin, or (**i**) Chx10 (arrowheads). (**j**) Quantification of the density of PRV^+^ neurons in grafts that express the phenotypic markers in **d-i**. The total density of PRV^+^ neurons per subject is shown to the right (n=19). (**k**) Correlation of graft volume vs. the density of PRV^+^ neurons in the graft for n=19 subjects. (**j**) Correlation of the total number of graft axons extending to 1000 μm caudal to the graft/host border vs. the density of PRV^+^ neurons in the graft for n=19 subjects. (**m**) Taking into account the abundance of each graft neuronal subtype, the percentage of subtype that is PRV^+^. All data are mean ± SEM. Scale bars = 100 μm (**b, c**), 10 μm (**d, e, f, g, h, i**). Figure was created with Biorender.com.

We also correlated the density of PRV^+^ GDNs to total graft volume and axon outgrowth. Surprisingly, we found that graft size was inversely correlated with the density of PRV^+^ neurons (p=0.0537, **Fig. 2k**). In contrast, we failed to observe any correlation between graft axon extension and graft infectivity with the PRV virus (**Fig. 2l**). This might be because polysynaptic relays do not require long-distance axon extension, but rather integrate through local synaptic connections. Finally, we quantified the mean abundances of graft neuronal phenotypes, then calculated the percentage of each phenotype that is PRV^+^, or the ‘connectivity index’ (**Fig. 2m**). It is important to note that the numbers of Chx10-immunoreactive cells are likely an underrepresentation of the actual counts of V2a interneurons in grafts, based on our previous findings that not all V2a GDNs express Chx10^35^. Through this approach, we found that of all the subtypes assessed, less than 5% of each subtype was labeled with PRV^+^. This indicates low overall connectivity but broad diversity in the types of neurons that spontaneously synaptically integrate into locomotor circuits.

### V0_C_ and V2a graft-derived neurons exhibit different axon projection patterns

NPC graft-derived neurons have been shown to extend large numbers of axons into the host spinal cord^9,11,44–46^. However, it is unknown whether distinct subtypes of GDNs may exhibit differential patterns of axon extension into the host spinal cord. To address this gap in knowledge, we next performed subtype-specific GDN projection mapping. We identified two subtypes of GDNs, cholinergic V0_C_ interneurons and glutamatergic V2a interneurons, as promising candidates for additional studies for the following reasons: (1) They are both premotor interneuron subtypes that modulate locomotor output in the intact spinal cord^47–51^, (2) both subtypes are present in mature NPC grafts^35^, and (3) we found that both of these GDN subtypes can synaptically integrate into locomotor circuits (**Fig. 2j**). To the best of our knowledge, all cholinergic GDNs are V0_C_ interneurons, not motor neurons. Phenotypically, these neurons have morphology consistent with small-diameter V0_C_ neurons. We have very rarely observed ChAT^+^ GDNs with MN morphology; namely, large soma size and extensive dendritic arborization.

To perform subtype-specific projection mapping of GDNs, we generated spinal cord NPC grafts using donor cells of the following genotypes: Syn1-cre (for Cre recombinase expression in all GDNs), ChAT-cre (Cre expression in V0_C_ GDNs), or Chx10-cre (Cre expression in V2a GDNs). Specificity of Cre-dependent gene expression in grafts was confirmed through transgenic crosses of Cre mice with Ai14 tdTomato reporter mice (**Fig. S2**). We transplanted Cre-expressing NPCs into sites of T12 SCI, and four weeks later injected AAV-SynTag (*AAV2-phSyn1(S)-FLEX-tdTomato-T2A-SynpEGFP-WPRE*) directly into grafts. This viral vector allows expression of cytoplasmic tdTomato and presynaptic (synaptophysin-fused) GFP in Cre-expressing neurons, allowing the visualization of graft-derived axons and their presynaptic terminals (**Fig. 3a-d**). We quantified caudal extension of tdTomato+ graft-derived axons from each graft type, and as expected, there were significantly more labeled axons in Syn1-cre grafts versus other graft types up to 2.5 mm caudal to the graft/host border (**Fig. 3e-f**). We next assessed the distributions of graft-derived GFP^+^ synaptic punctae in transverse sections of host lumbar spinal cord (**Fig. 3g**). We generated density maps of GFP^+^ punctae at spinal levels L3-L6 to visualize areas of greatest synaptic density for each graft type (**Fig. 3h**). We quantified the percentages of total graft-derived synaptic punctae in each Rexed laminae at spinal level L3, and found that most of the terminals were located in laminae V and VII-VIII (**Fig. 3h, i**). As expected, Syn1-cre grafts exhibited the greatest numbers of synaptic punctae in the lumbar spinal cord compared with ChAT-cre and Chx10-cre grafts (**Fig. 3h**). Notably, presynaptic density for cholinergic GDNs was significantly enriched in lamina V, and presynaptic density for V2a GDNs was significantly enriched in laminae VII-VIII (**Fig. 3i**). This is consistent with the projection patterns of these neuronal subtypes in the native spinal cord: V2a interneurons synapse onto targets in laminae VII-VIII including V0_D_ and V0_V_ populations^47,52^, and V0_C_ primarily project to the ventral horn and intermediate gray matter^48^.

**Figure 3.**
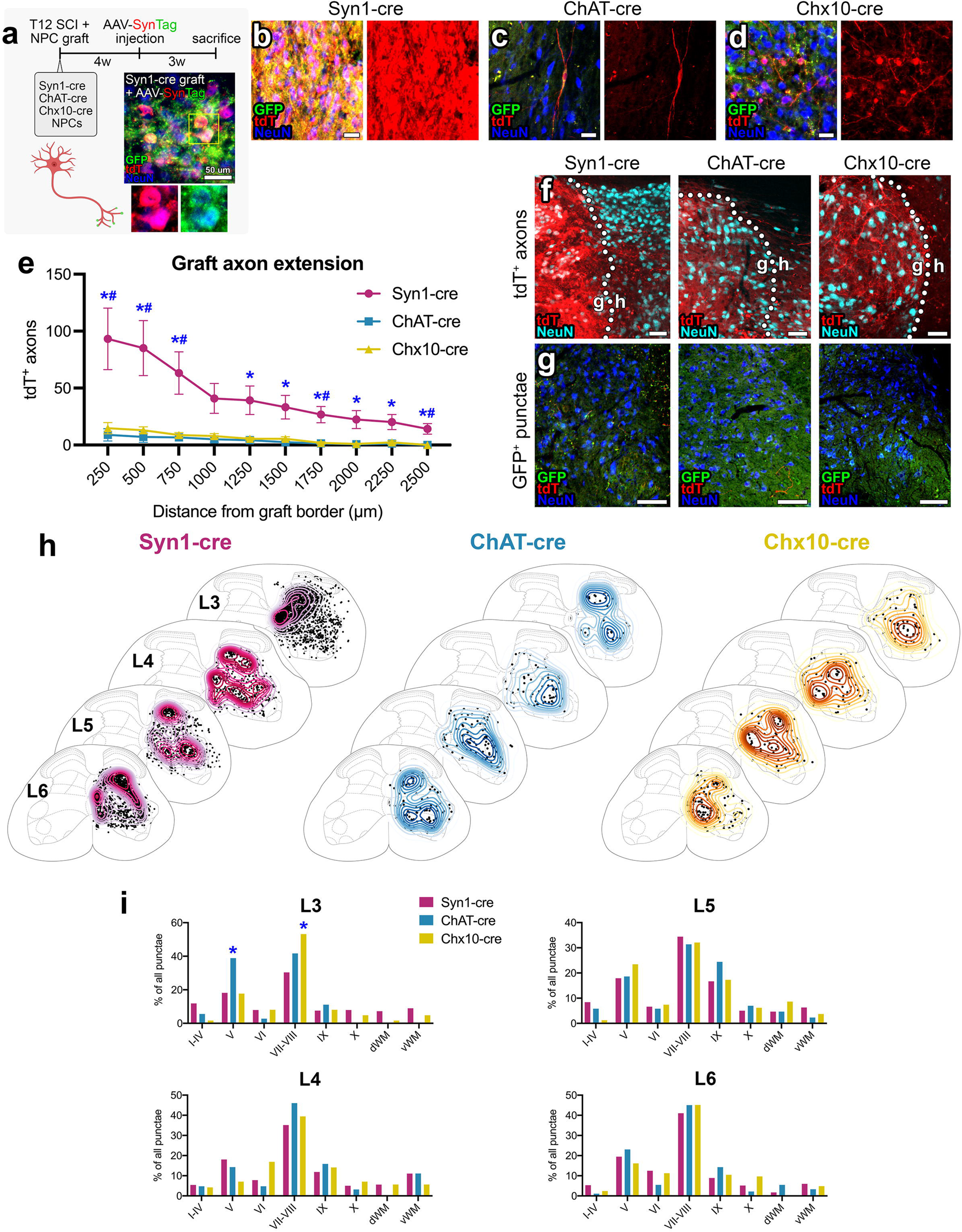
Graft-derived neurons form synaptic contacts in the intermediate and ventral laminae of the host lumbar spinal cord. (**a**) Experimental strategy. Dorsal column lesion SCI was administered at spinal level T12, followed by immediate transplantation of either Syn1-cre, ChAT-cre, or Chx10-cre NPCs into the lesion cavity. Four weeks later, grafts were injected with Cre-dependent AAV-SynTag (cytoplasmic tdTomato + synaptophysin-EGFP fusion). Three weeks later, animals were sacrificed for histological analysis. The immunofluorescent panel is a high-magnification image within a Syn1-cre graft labeled with AAV-SynTag. (**b-d**) Representative images of grafts with tdTomato and GFP expression in (**b**) all neurons, (**c**) V0_C_ graft neurons, or (**d**) V2a graft neurons. (**e**) Quantification of total outgrowth of tdTomato^+^ axons from each graft type at 250-μm intervals caudal to the graft/host border. *p<0.05 for Syn1-cre vs. ChAT-cre; #p<0.05 for Syn1-cre vs. Chx10-cre. (**f, g**) Representative images of (**f**) tdTomato expression in graft-derived axons at the graft (g) / host (h) border, and (**g**) GFP expression in graft-derived synaptic terminals within the host lumbar (L3-L6) spinal cord gray matter. (**h**) Density maps of synaptic punctae for each graft type at lumbar spinal cord levels L3, L4, L5, and L6. (**i**) Density of synaptic punctae from each graft type within the L3 host spinal cord, by laminae. *p<0.002 significantly higher than random chance by Monte Carlo approach. All data are mean ± SEM. Scale bars = 100 μm (**f, g**), 50 μm (**b, c, d**). Figure was created with Biorender.com.

### V0_C_ and V2a graft neurons can functionally integrate into spinal locomotor circuits

Through the above experiments, we found that GDNs including V0_C_ and V2a interneurons can synaptically integrate into spinal locomotor circuits. We next sought to determine whether GDNs can functionally integrate into these circuits. To explore this possibility, we generated mouse embryos with Cre-dependent hM3Dq (Gq-DREADD) expression in all GDNs (Syn1-cre::hM3Dq) and transplanted spinal cord NPCs into sites of SCI in wild-type mice (**Fig. 4a**). To validate graft hM3Dq activity, at 4 weeks post-transplantation we injected animals intraperitoneally with the DREADD agonist, clozapine-*N*-oxide (CNO), and sacrificed subjects for histology 45 minutes later. Immunolabeling for the immediate early gene c-Fos, a reporter of neuronal activity^53,54^, revealed that activation of Syn1-cre::hM3Dq grafts with CNO was effective in increasing GDN activity (**Fig. 4b-c**). In addition, some of the animals were used for electrophysiology experiments (**Fig. 4d-e**). Four weeks after transplantation of either Syn1-cre::hM3Dq NPCs or GFP^+^ NPCs (control), animals were anesthetized and 32-channel silicon microelectrode arrays were lowered directly into grafts. Baseline recording was performed for 10 minutes, then CNO was injected and recordings continued for 45 minutes (**Fig. 4d**). Following CNO injection, there was a sharp and significant rise in firing rates of neurons within h3MDq grafts, but not control GFP grafts (**Fig. 4e**). Collectively, these data validate that activity of GDNs can be modulated via a chemogenetic approach.

**Figure 4.**
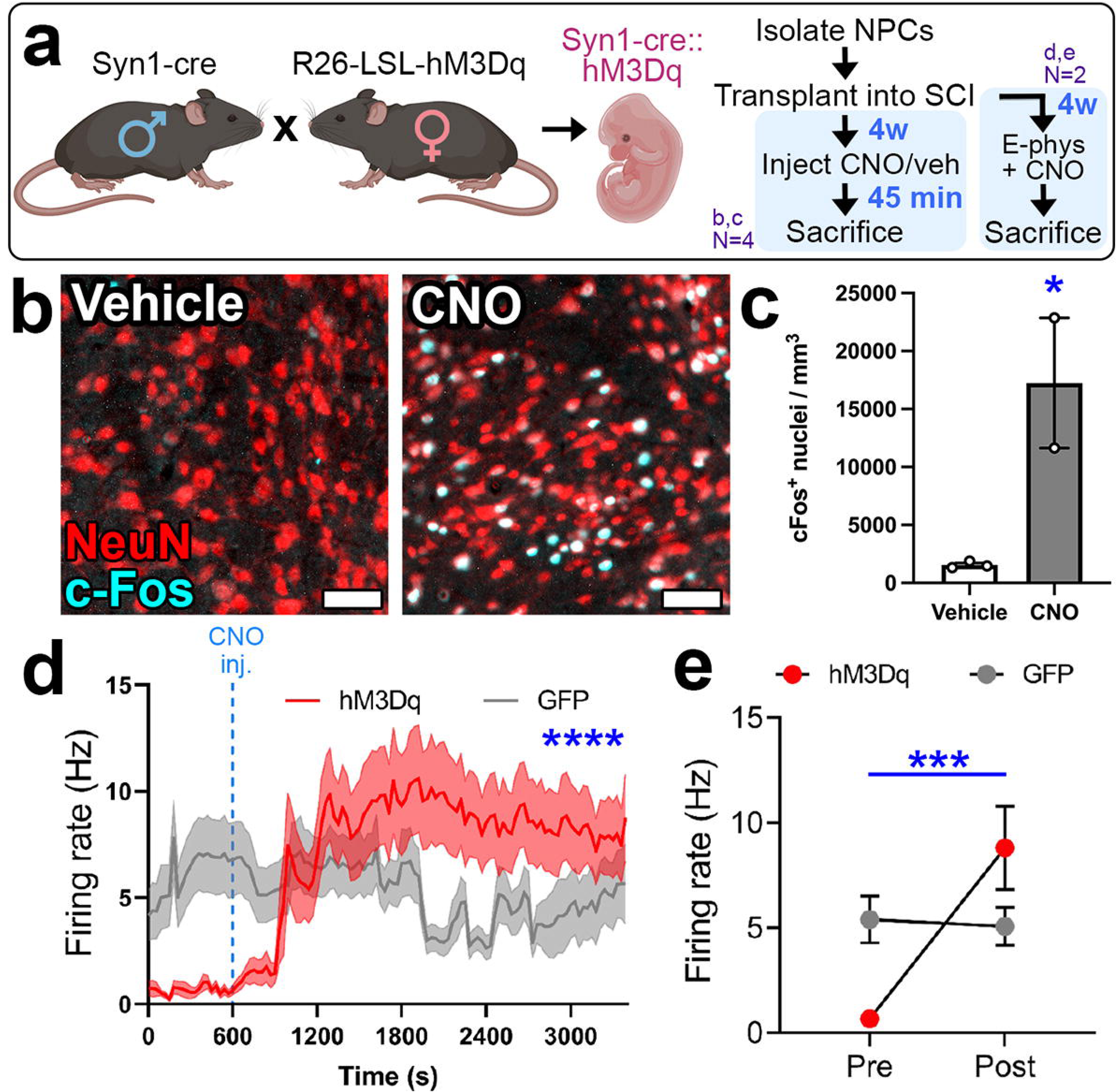
Validation of hM3Dq expression and function in Syn1-cre::hM3Dq NPC grafts. (**a**) Approach for generating Gq-DREADD expression in graft-derived neurons. Syn1-cre males and cre-inducible hM3Dq females were paired to generate Syn1-cre::hM3Dq embryos for donor NPCs. NPCs were transplanted into sites of cervical SCI; 4 weeks later, in vivo electrophysiological recordings and c-Fos expression analysis were performed. (**b, c**) c-Fos expression in Syn1-cre::hM3Dq grafts 45 minutes after i.p. injection of either vehicle or clozapine-N-oxide (CNO). *p<0.05 by unpaired, two-tailed t test. (**d**) Time series of neuron firing rates in control GFP graft (n=36 neurons) or Syn1-cre::hM3Dq graft (n=47 neurons) *in vivo* over 55 min (CNO injection at 10 min). ****p<0.0001 time x graft type main effect by two-way RM ANOVA. (**e**) Mean firing rates pre- and post-CNO injections. ***p=0.0004 time x graft type main effect by two-way RM ANOVA. All data are mean ± SEM. Scale bars = 50 μm (**b**). Figure was created with Biorender.com.

We next assessed whether increasing activity of GDNs could result in hindlimb muscle activation. To explore this possibility, we generated NPC grafts with Cre-dependent hM3Dq expression in either all GDNs (Syn1-cre::hM3Dq), cholinergic GDNs (ChAT-cre::hM3Dq), or V2a graft neurons (Chx10-cre::hM3Dq; **Fig. 5a**). At 9 weeks post-transplantation into T12 contusion SCI, we assessed open locomotor scores both prior to, and immediately following, i.p. injection of CNO. We did not detect any significant differences in pre-CNO vs. post-CNO locomotor scores (**Fig. 5b**). As such, we opted to more subtly probe behavior using motion sequencing (MoSeq)^55–60^. MoSeq is a data-driven, unsupervised machine learning approach designed to systematically identify and classify behavioral structure in freely moving mice. MoSeq captures sub-second behaviors and segments them into discrete, recurring units known as “syllables” (rear, groom, locomotion, etc.). MoSeq can also provide insight into the sequential organization of syllables (“bigram probability”) in complex motor behavior^55–58^. Following injection of CNO, we observed significant differences in behavioral patterns between the control and Syn1-cre::hM3Dq groups across multiple syllable categories including grooming, locomotion, and rearing (**Fig. 5c**), while the ChAT-cre::hM3Dq and Chx10-cre::hM3Dq groups exhibited fewer changes. Additionally, we identified conserved changes in bigram probability across all three grafted groups, characterized by a down-regulation in pause/freeze-to-groom transitions and an upregulation in pause/freeze-to-rear sequences (**Fig. 5d)**. Together, these findings indicate subtle differences in treatment effects on gross motor behaviors, revealing graft-driven shifts in behavioral state dynamics.

**Figure 5.**
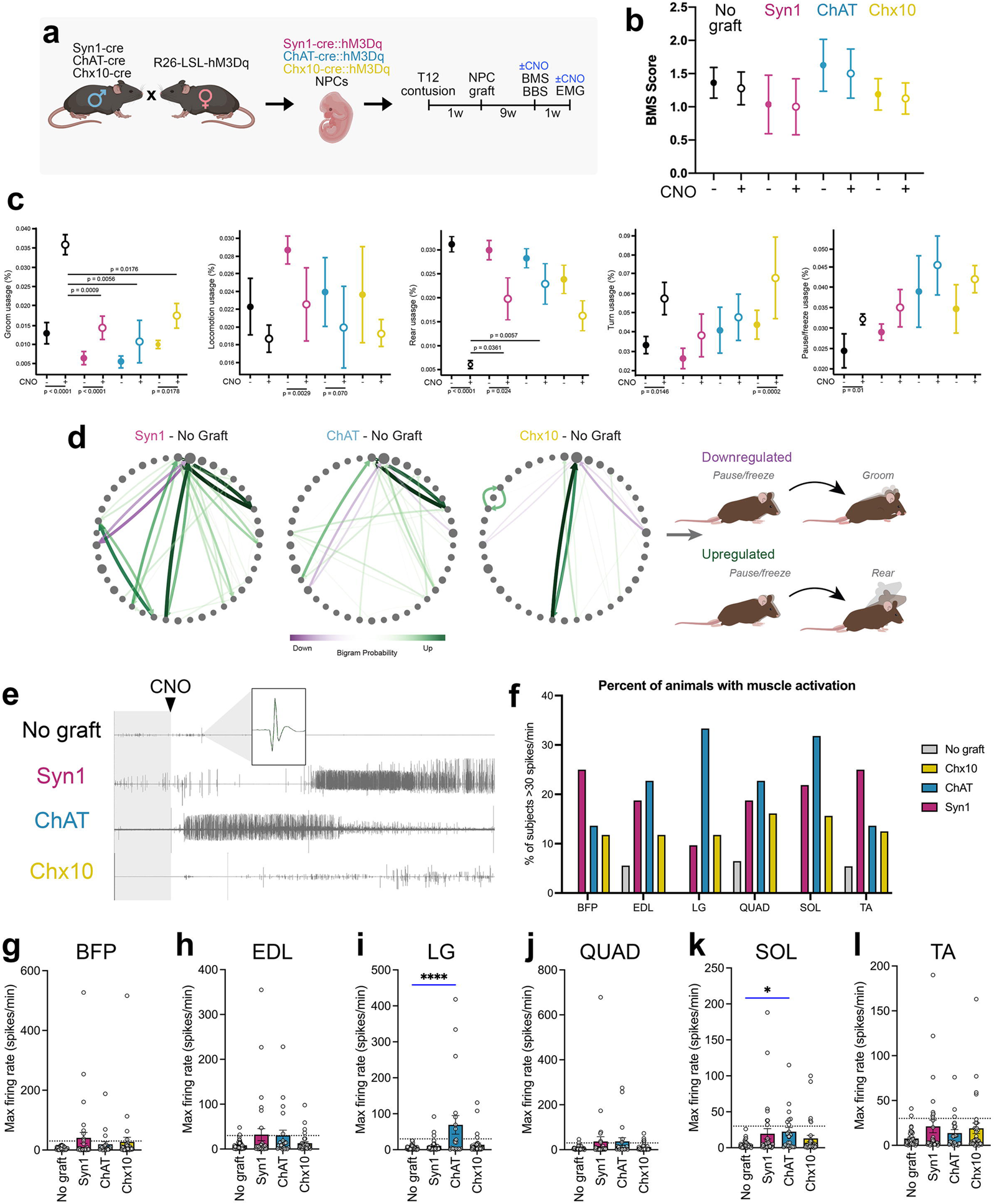
Activation of graft-derived neurons is sufficient to elicit hindlimb muscle activation. **(a)** Experimental strategy and timeline. Adult mice received transplantation of Syn1-cre::hM3Dq, ChAT-cre::hM3Dq, or Chx10-cre::hM3Dq NPCs into sites of T12 contusion. Nine weeks later, open field locomotor (Basso Mouse Scale) scoring and terminal electromyography recordings were performed pre- and post-CNO injection. (**b**) BMS scores for animals with each graft type pre- and post-CNO injection in the same testing session. (**c**) Subsecond behavioral patterns (“syllables”) identified through motion sequencing exhibit significant changes following CNO injection. (**d**) Behavioral statemaps, normalized to the control group, reveal conserved upregulation and downregulation of specific syllable sequences across experimental groups. (**e**) Example EMG traces for animals with each graft type. Time of CNO injection is indicated with an arrowhead. A representative waveform is shown in the inset. (**f**) The percent of animals with CNO-induced activation of the lateral gastrocnemius muscle, one of 6 muscles recorded in this study. *p<0.05, ****p<0.0001 vs. no graft by one-way ANOVA + Dunnett’s multiple comparisons test. Figure was created with Biorender.com. (**g-l**) Maximum muscle activation rate for individual muscle groups (BFP: biceps femoris posterior; EDL: extensor digitorum longus; LG: lateral gastrocnemius; QUAD: quadriceps; SOL: soleus; TA: tibialis anterior) during the recording period following CNO delivery. Each data point represents an individual subject.

At 10 weeks post-transplantation, we performed electromyographical recordings of 6 hindlimb muscles (biceps femoris posterior, extensor digitorum longus, lateral gastrocnemius, quadriceps, soleus, and tibialis anterior) in the left and right hindlimbs of anesthetized animals. Recordings were obtained for 15 minutes prior to CNO injection (baseline) then for 75 minutes following i.p. CNO injection. Prior to CNO injection, baseline activation rates were typically below 10 spikes/min for all muscles (**Fig. S3**). Example single-muscle recordings are shown in **Fig. 5e**. Following CNO administration, we found that some muscles in animals grafted with Syn1-cre::hM3Dq, ChAT-cre::hM3Dq, and Chx10-cre::hM3Dq NPCs exhibited increased activity compared to baseline. Root mean square data is shown in **Fig. S4**. The percentages of subjects with muscle activation are presented in **Fig. 5f**. To determine the effects of GDN activation on muscle activity, we quantified the maximum activity (spikes per minute) for each muscle within the 75-minute period following CNO administration (**Fig. 5g-l**). In most (205 of 211; 97.2%) cases, individual muscles of ungrafted (control) animals did not exceed 30 spikes/min throughout the entire recording period. This slight increase over baseline activity may indicate slight off-target effects of CNO on muscle activation. We therefore set a threshold of 30 spikes/minute to qualify as muscle “activation” above baseline. Using this criterion, a subset of animals (“responders”) in each of the three grafted groups exhibited heightened muscle activity following CNO administration for each of the muscles recorded, in some cases firing at several hundred spikes/min (**Fig. 5g**). The observation that many animals in each of the grafted groups failed to exhibit increased muscle activity after CNO indicates that not all grafted animals had sufficient numbers of graft-to-motor circuit connections to drive activity. Together, these findings indicate that activation of graft-derived neurons is sufficient to drive activity of locomotor circuits resulting in hindlimb muscle activation.

## Discussion

A considerable number of experimental animal studies have demonstrated therapeutic effects of transplanted neural stem- and progenitor cells on voluntary motor function, but mechanisms of graft-mediated functional recovery after SCI remain unclear. Recently, Zholudeva and colleagues demonstrated connectivity of grafted human V2a spinal interneurons (V2a SpINs) with spinal respiratory circuits following high cervical SCI^43^. Optogenetic manipulations demonstrated that neural relays between host and graft neurons supported recovery of diaphragm function, providing novel evidence that host/graft synaptic connectivity can improve functional outcomes after SCI. Here, we have demonstrated that graft-derived neurons including but not limited to V0_C_ and V2a subtypes are synaptically poised to modulate hindlimb muscle activity. Graft-derived neurons spontaneously establish synaptic connections with host locomotor circuitry, and chemogenetic activation of graft neurons was sufficient in a subset of mice to elicit hindlimb muscle activation. One limitation of our study is that we did not investigate the other half of the equation to drive voluntary locomotor function: host-to-graft inputs. Mouse locomotor recovery after SCI has been shown to be driven by spontaneous or experimentally manipulated plasticity in supraspinal systems such as cortical and brainstem centers^61–65^ as well as plasticity in the propriospinal system^66–71^. It is possible that these motor pathways could functionally integrate with grafts to drive additional recovery. Although outside the scope of the current study, this is a critical topic for future investigation.

The cell type-specific nature of graft-to-host connections has not previously been investigated in NPC transplantation studies. We previously showed that regenerating host axons can discriminate between phenotypically appropriate and inappropriate targets within NPC grafts^7^, highlighting the ability of adult axons to ‘remember’ their correct postsynaptic targets after injury. Our present data suggest that graft-derived neurons may also be able to identify their own phenotypically correct postsynaptic targets within the host spinal cord, as termination patterns within distinct spinal laminae differed between GDN subtype. Collectively, these observations underscore the possibility that NPC transplantation might accurately reconstruct damaged motor circuits after SCI. Future work is needed to characterize the precise nature of the synaptic connections that are spontaneously established between host and graft. In the chemogenetic component of our study, we found that only a subset of animals exhibited increased muscle activation upon graft activation. Even in animals that expressed the excitatory DREADD in all graft neurons, only about 25% of subjects responded to graft activation on EMG recordings. While these data highlight the sufficiency of graft-to-host synapses to drive muscle activity, they also demonstrate that most animals receiving NPC transplantation do not establish enough synaptic connections to result in functional changes. Importantly, we observed that the vast majority of Syn1-GFP labeling was found within the grafts themselves, suggesting that newly grafted neurons preferentially synapse with each other rather than host neuronal targets. This underscores the need for additional efforts to drive graft-to-host, rather than graft-to-graft, synaptic connections.

In the current study as well as in a recently published study by our lab^35^, we failed to observe any improvements in locomotor function associated with NPC transplantation in a mouse thoracic contusion SCI model. However, we found that chemogenetic activation of graft-derived neurons shifted behavioral states away from static-like behaviors toward more dynamic behavioral patterns. Our conservative interpretation is that while NPC transplantation may influence motor behavior in subtle ways, it is insufficient on its own to drive meaningful locomotor recovery in a clinically relevant thoracic spinal cord contusion SCI mouse model. This appears to contradict multiple reports of graft-associated improvements in locomotion following SCI published by other groups^5,18–20,22,45,72–74^. What underlies this discrepancy? It is possible that there are key differences in the experimental approaches used between different research groups. We propose that hidden factors in what defines a graft’s “success”, such as biological sex^9^ or graft cellular composition^35^, may represent key sources of variability in outcomes for grafting studies, not only in SCI, but other models of neurological disease and trauma such as traumatic brain injury, stroke, and epilepsy. In the current study, we observed that the percentage of graft neurons that are synaptically integrated with host locomotor circuits is quite low, representing less than 1% of all graft neurons. This highlights the need for strategies to enhance host-graft connectivity, optimize graft composition, and/or combine transplantation with other growth-promoting treatments in order achieve biologically meaningful improvements in locomotor function.

## Materials and Methods

### Ethics statement

Animal studies were performed in compliance with the *NIH Guidelines for Animal Care and Use of Laboratory Animals*. All experiments utilizing animals were approved by the Texas A&M University Institutional Animal Care and Use Committee. All efforts were made to minimize pain and distress.

### Animals

A total of 247 experimental mice were used for this study, plus 120 additional mice for embryo generation. Genotypes of mice included C57BL/6J (#000664), GFP [C57BL/6-Tg(CAG-EGFP)131Osb/LeySopJ; Jax #006567], Ai14 [B6;129S6-Gt(ROSA)26Sortm14(CAG-tdTomato)Hze/J, Jax #007908], CAG-LSL-hM3Dq [B6N;129-Tg(CAG-CHRM3*,-mCitrine)1Ute/J, Jax #026220), Syn1-cre (B6.Cg-Tg(Syn1-cre)671Jxm/J, Jax #003966), ChAT-cre (B6;129S6-Chattm2(cre)Lowl/J, Jax #006410), and Chx10-cre [Tg(Vsx2-Cre)TC9Gsat/Mmucd, MMMRRC #36672]. Equal numbers of adult (8-12 week old) animals of both sexes were used for all experiments. Animals had free access to food and water throughout the study and were group-housed in standard Plexiglas cages on a 12-hour light / 12-hour dark cycle (light cycle = 6:00 am – 6:00 pm). Animal housing facilities had ambient temperature between 20-23°C and 30-70% humidity. Thirty-four animals were removed from the study due to premature mortality and nineteen subjects were removed due to poor graft survival.

### Pre- and post-operative surgical procedures

All surgeries were performed under anesthesia using a combination of ketamine (25 mg/kg), xylazine (5.8 mg/kg), acepromazine (0.25 mg/kg), and inhaled isoflurane (0.5%-1.5%). Following induction of anesthesia, surgical sites were shaved and sterilized with alternating scrubs of betadine and 70% ethanol. Following surgery, antibiotic powder (Neo-Predef, Covetrus) was applied to the surgical site prior to closure of incised skin by stainless steel wound clips. Post-operative care included subcutaneous injection of banamine (0.05 mg/kg) and ampicillin (0.05 mg/kg) in lactated Ringer’s solution once daily for 72 hours. Immediately after surgery, animals were returned to their cages and remained half on/half off heating pads for 72 hours after surgery. Daily health checks, bladder care, and body weight measurements were performed throughout the duration of these studies.

### Virus injections

#### AAV helper virus and modified rabies virus

A subset of animals (**Fig. 1**) received injection of AAV helper virus (AAV2/1-flex-N2cG-histHA at 2.4x10^10^ gc/mL; AAVDJ-flex-TVA-mCherry at 1x10^11^ gc/mL; both purchased from Dr. Andrew Murray, University College London) into the L2-L5 spinal segments. Injections were made using the Nanoject III (Drummond). A total of 12 injections of 300 nL each were delivered across the following coordinates across L2-L5 spinal levels: medial/lateral: ± 0.3mm, and dorsal/ventral: -0.5 mm to -0.7 mm. The same subset of animals received injections of modified rabies virus (RABV-N2CdeltaG-TdTomato at 10^7^ gc/mL; purchased from Dr. Andrew Murray, University College London) 12 weeks later. A total of 12 injections of 300 nL each were delivered into the same coordinates as above.

#### Pseudorabies virus

A subset of animals (**Fig. 2**) received injections of pseudorabies virus (PRV-614 at 6x10^7^ gc/mL; purchased from the NIH Virus Center) into the sciatic nerves. A total of 6 injections of 300 nL each were made into the sciatic nerve.

#### AAV-SynTag

A subset of animals (**Fig. 3**) received injections of AAV-SynTag (AAV2-phSyn1(S)-FLEX-tdTomato-T2A-SynpEGFP-WPRE at 10^11^-10^12^ gc/mL; Addgene #51509) into NPC grafts. A total of 16 injections of 250 nL each were delivered across the following coordinates at the T12 spinal cord level: medial/lateral ± 0.3 mm, dorsal/ventral -0.5 mm and -0.7 mm.

### Spinal cord injury

#### Contusions

A laminectomy at vertebral level T12 was performed. After laminectomy, the T11 and T13 foramen were then clamped to stabilize the spinal cord prior to contusion. A contusion to spinal level T12 was then delivered with the Infinite Horizon Impactor (IH-0400, Precision Systems and Instrumentation, Fairfax, VA; 60 kydnes, 1 second dwell), using a 1.3-mm impactor tip.

#### Dorsal column lesions

Dorsal column lesions were performed at vertebral level T12. Following laminectomy, a tungsten wire knife with an extruded diameter of 1.0-1.5 mm (McHugh Milieux, Downers Grove, IL) was centered above the spinal cord midline, retracted, and inserted to a depth of 0.8 mm below the dorsal spinal cord surface. The knife was then extruded and raised to transect the spinal cord dorsal column. The completeness of the injury was verified upon visual inspection.

### Neural progenitor cell isolation and transplantation

#### NPC isolation

Mouse embryos were generated through timed mating^9^. Adult wild-type female mice received intraperitoneal injections of luteinizing hormone-releasing hormone (5 I.U.; Sigma-Aldrich) before 10:00am. Four days later, injected females were paired with homozygous GFP adult male mice overnight. On the day of transplantation, embryonic day 12.5 (E12.5) embryos were harvested. Embryonic spinal cords were collected in ice-cold Hank’s Balanced Salt Solution (HBSS) then digested in 0.125% trypsin for 12 minutes at 37°C. A cocktail of 10% fetal bovine serum in Dulbecco’s Modified Eagle Medium was then added to neutralize the reaction before being centrifuged at 600 RCF for 2.5 minutes. The supernatant was then removed and the tissue was then gently resuspended in Neurobasal Medium + 2% B27 Supplement. Cell viability was assessed by trypan blue exclusion. Cells were stored on ice in NBM + B27 until transplantation.

#### NPC transplantation

For transplantation into sites of spinal cord contusion, the lesion site was re-exposed at 1 week post-SCI. NPCs were resuspended in a fibrin matrix to a concentration of 300,000 viable cells/µL. A 4 µL total volume of cells was injected into the lesion site at a depth of 0.5 mm - 0.8 mm across 6 injection sites via pulled glass micropipette using a PicoSpritzer II (General Valve, Inc. Fairfield, NJ), over a period of 3-5 minutes. Subjects were randomly assigned to receive NPCs or vehicle (fibrin matrix only). For transplantation into sites of dorsal column lesion, 2 µL of NPCs at a concentration of 500,000 viable cells/µL suspended in HBSS were injected into the injury cavity at a depth of 0.7 mm via a pulled glass micropipette using a PicoSpritzer II.

### Open field locomotor assessments

Functional recovery of hindlimbs was assessed using the Basso Mouse Scale for Locomotion^75^. Animals were tested in rooms where their home cages were located and allowed to explore the testing arena prior to scoring. Animals were allowed to roam freely in a 14 inch by 30 inch plexiglass box for four minutes under the supervision of two scorers. Plexiglass box was wiped down with 0.3% acetic acid between animals. Baseline scores were collected prior to surgery, and all mice exhibited baseline BMS scores of 9. Following SCI, BMS scores were obtained on 1-, 3-, 6-, and 14 DPI, then once weekly until 56 DPI. At each time point, scores for both hindlimbs were averaged to produce a final score.

### Motion Sequencing

Motion sequencing was carried out as previously described^55–60^.

#### Data acquisition

Prior to data acquisition, mice were allowed to habituate to the behavior room for 30 minutes. Raw depth and color video data was acquired using a custom C# program, kinect2-nidaq (v0.2.4alpha), installed on a computer connected to a Microsoft Kinect v2 camera which was positioned above a 17-inch diameter circular arena resulting in a top-down view of the arena interior. Individual mice were gently placed in the area and allowed to behave naturally over a 20-minute period while their movements were recorded. Depth frames, encoded as 16-bit unsigned integers encoding the distance from the camera sensor, were captured at 30Hz with a resolution of 512 x 424 px. Data was saved in a binary format and transferred to another computer system for later processing.

#### Data preprocessing

Raw depth data was processed as described previously^56^. Briefly, we used a custom developed deep learning network, based on the popular mask-R-CNN architecture, to identify and segment mice from the raw depth maps, and produce output files compatible with the ecosystem of MoSeq software.

#### Modelling

Preprocessed data was submitted to principal component analysis to transform depth data into a 10-dimentional time series of PCA scores, which describe the animals’ movement dynamics. Optimal kappa parameters were discovered via grid-search with a target of minimizing the difference between the modelled mean syllable duration and the that of a mode-free changepoint analysis. A family of ten robust models was trained for 1,000 iterations using the optimal kappa and we chose the model with median log-likelihood to take forward for further analysis.

#### Posthoc analyses

Behaviors, henceforth referred to as “syllables,” were computationally identified and then categorized using the following semantic labels: locomotion, groom, rear, pause/freeze, and turn. Syllables were quantified based on their frequency of occurrence (“usage”). To eliminate erroneous data points, a bootstrap median was computed, and Tukey’s fences were applied using the median absolute deviation to identify and remove outliers. Statistical analyses included one-way ANOVA with Tukey’s HSD to assess differences between groups within conditions (CNO-, CNO+). Additionally, a linear mixed effects (LME) model was used to evaluate within-group differences across conditions. In the LME, usage was the dependent variable, group (CNO-as the baseline) was the fixed effect, and animal was included as a random effect to account for missing data, uneven sample sizes, and repeated measures common in SCI studies.

### Electrophysiology

A total of 5 mice underwent acute spinal cord electrophysiological recordings to confirm CNO-mediated activation of hM3Dq ^+^ grafts. Uninjured mice were stereotaxically injected with NPCs of either the Syn1-Cre::hM3Dq or GFP genotype into the dorsal columns at the T12 spinal level. Four weeks later, mice were placed into a stereotaxic frame and the T12 spinal cord was re-exposed. The location of the graft was verified upon visual inspection. A 32-channel silicon microelectrode array (Buzsaki-A32, NeuroNexus, Ann Arbor, MI) comprised of four 8-channel probes was used for extracellular recordings and the pre-amplifier was directly grounded to the stereotaxic frame. In short, after the silicon probe was inserted into the graft, it was slowly advanced until good signal was visually observed across most channels. The signal was then allowed to stabilize for ∼30 minutes before recordings began. Ten minutes after recordings began, mice received i.p. injections of the selective hM3Dq agonist, CNO (5 mg/kg in DMSO). Recordings continued for 1-hour post injection to determine if hM3DQ activation via CNO increased the firing rate of recorded spinal units. After the experiment, mice were euthanized with an anesthetic cocktail overdose and transcardially perfused with physiological saline followed by 4% paraformaldehyde; spinal cord tissue was collected for histology. Extracellular single-unit activity was acquired by a multichannel neurophysiological recording system (OmniPlex, Plexon, Dallas, TX) as previously described^76^. Wideband signals were acquired at 40-kHz on each channel, amplified (2000x), and saved offline for further analysis. Only waveforms that exceeded a threshold of 3 standard deviations below baseline noise were selected for unit sorting. After bandpass filtering (600-6000 Hz), waveforms were manually sorted using the first two principal components (Offline Sorter, Plexon). Only well-isolated units were considered for further analysis. Using NeuroExplorer (Plexon), the average firing rate of sorted units was analyzed from a period of 10 min pre-injection to 45 minutes post-injection using 30 second bins.

### Electromyography

In Figure 5 experiments, mice underwent electromyographical recordings at 10 weeks post-NPC transplantation. Before electromyography recordings began, 12 handmade recording electrodes were placed in bilateral biceps femoris posterior, quadriceps (QUAD), gastrocnemius lateralis, soleus, extensor digitorum longus, and tibialis anterior muscles of anesthetized mice. The ground electrode was placed subcutaneously on the dorsal aspect of the mouse between the left and right legs. Muscle activity was recorded first received to a Model 3500 16-Channel Extracellular Differential Amplifier (A-M Systems). During amplification, muscle activity was filtered through a high pass filter=30, low pass filter=1000, gain=500, and inclusion of notch filtering. Data was then digitized by a Micro3 1401 digitizer (A-M Systems) that was then presented as 12-channel waveform data in Spike2 software version 10. Post-recording, muscle activity was filtered through a 100Hz low pass filtered. Muscle activity was analyzed through the quantification of spikes that hit at least a ±5 standard deviation of the mean activity of each electrode channel. Root mean square (RMS) analysis was performed through the Spike2 software (Cambridge Electronic Design).

### Euthanasia and tissue processing

Animals were euthanized by an anesthetic cocktail overdose. After overdose, animals were transcardially perfused with 50 mL of physiological saline followed by 50 mL of 4% paraformaldehyde (PFA) in 0.1M phosphate buffer. Spinal columns were post-fixed in 4% PFA overnight at 4°C. After post-fixation, spinal cords were then stored in 30% sucrose at 4°C for at least 72 hours. A 1-cm length of spinal cord centered around T12 was removed and frozen in Tissue-Tek OCT compound (VWR) on dry ice. Spinal cord tissue was cryosectioned in the sagittal plane to a thickness of 20µm. Sections were stored at 4°C.

### Immunohistochemistry

As previously described, tissue was first washed in tris buffered saline (TBS) three times for 10 minutes each before a one-hour incubation in blocking solution containing TBS, 0.25% Triton-X-100 (Sigma-Aldrich), and 5% donkey serum (Lampire Biological Laboratories). Tissue was then transferred into primary antibody solution for overnight incubation at 4°C. The next day, tissue was washed again in TBS then incubated in blocking solution containing AlexaFluor-conjugated secondary antibodies (Jackson ImmunoResearch). Primary and secondary antibodies used for immunohistochemistry are listed in **Table S1**. Tissue was then incubated in DAPI solution (5 μg/mL, Sigma-Aldrich) before being mounted onto slides and coverslipped using Mowiol mounting medium.

### Optical clearing and lightsheet microscopy

Spinal cord samples were obtained following transcardial perfusion with ice-cold phosphate-buffered saline (PBS) followed by 4% PFA solution. The spinal column was removed and post-fixed in a 4% PFA solution for 24 hours before being washed with PBS. Spinal cords were removed from the spinal column and cleared using a modified iDISCO^+^ protocol^77,78^. Briefly, samples were dehydrated in a methanol/H_2_O gradient series and delipidated in a 2:1 dichloromethane:methanol solution overnight. Samples were then rehydrated with a reverse methanol/H_2_O gradient before being permeabilized with a solution of PBS, 0.2% triton-X, glycine (2.3% w/v) and dimethyl sulfoxide (20% v/v) at 37°C overnight. Spinal cords then underwent blocking with 5% normal donkey serum in PBS containing 0.2% Tween-20, 0.02% sodium azide, and 0.01% (w/v) heparin (PTwH solution) overnight at 37°C before immunofluorescent staining with goat anti-Choline Acetyltransferase (1:400, Genetex GTX8275) and rabbit anti-RFP (1:400, Rockland 600-401-379) in PTwH for 2 days at 37°C. Primary antibodies were washed off with PTwH and spinal cords were then incubated with Alexa Fluor 647 anti-goat (1:200, Invitrogen A21447) and Alexa Fluor 555 anti-rabbit (1:200, Invitrogen A32794) secondary antibodies in PTwH for 2 days at 37°C. After two days, spinal cords were washed with PTwH, mounted in 2% low-melt agarose, dehydrated in a methanol/H_2_O series, and incubated before a final delipidation in 66% DCM/33% methanol for 3 hrs. Residual methanol was removed with two washes in 100% DCM for 15 min each and samples were transferred to 5 mL of room temperature ethyl cinnamate for 24 hours for refractive index matching. Images were captured using a Zeiss Lightsheet Z1 Fluorescent microscope, and images were processed using Imaris software (Bitplane).

### Image analysis

All image analysis was performed in a blinded fashion using the FIJI software. Regions of interest (ROIs) were drawn around grafts using fluorescent signal as a visual guide. All colocalization quantification was performed manually.

#### Axon outgrowth quantification

tdTomato fluorescence was overexposed to identify axons distal to the graft site. Using FIJI, graft ROIs were translated 250 µm in the rostral direction for 2 mm to identify the total number of axons crossing the leading edge of the ROI. Data is represented as the total graft axon outgrowth, normalized to graft volume.

#### Synaptic punctae density quantification

A Chi-Square test was first performed to determine whether cell counts varied among regions and graph types. Because counts in some regions were less than five, p-values were calculated via simulation^79^. A Monte Carlo approach was then used to assess significance of differences in individual cell counts across regions and graph types. We first calculated the proportion of cells observed in each region by pooling all graph types (e.g., for L3: 0.06, 0.25, 0.06, 0.42, 0.09, 0.04, 0.03, 0.05). We then used these proportions to create simulated datasets where the region of observation for cells in each graph type was sampled based on the above proportions. For each graph type, the simulated dataset’s sample size was equal to the empirical data. We repeated this process 10000 times to generate null distributions and used these distributions to calculate empirical p-values (two-tailed) for each observation. Based on a Bonferroni correction, we considered results significant at an alpha level of 0.002

### Statistical analysis

GraphPad Prism 8 (GraphPad Software, Inc.; La Jolla, CA) and custom software were used to perform statistical analysis. All data are presented as mean ± SEM. Statistical significance was defined as pL0.05. Details of statistical analyses are provided in **Table S2**.

## Author Contributions

A.T. contributed to study design, performed experiments, analyzed data, and wrote the manuscript.

A.B. contributed to data analysis, animal care, behavior testing, tissue processing, cryosectioning and immunohistochemistry.

J.T.E. contributed to MoSeq data processing, modelling, and analysis.

J.K.T. contributed to MoSeq data processing, modelling, and analysis.

K.V. contributed to data analysis, animal care, behavior testing, microscopy, and immunohistochemistry.

H.T. contributed to data analysis, animal care, behavior testing, tissue processing, cryosectioning, and immunohistochemistry.

A.T. contributed to data analysis, tissue processing, cryosectioning, and immunohistochemistry.

J.M. contributed to data analysis, behavior testing, tissue processing, cryosectioning, and immunohistochemistry.

B.S. contributed to data analysis, tissue processing, cryosectioning, and immunohistochemistry

T.G. contributed to data analysis.

A.S. contributed to data analysis.

A.P. contributed to data analysis.

S.N. contributed to data analysis.

M.P. contributed to behavioral testing and tissue processing.

S.L. contributed to behavioral testing and tissue processing.

J.J. contributed to behavioral testing, tissue processing, cryosectioning, and immunohistochemistry

M.T. contributed to contributed to electrophysiological recordings.

F.L.J. contributed to tissue clearing and lightsheet microscopy.

M.A. contributed to animal surgeries and animal husbandry.

A.F.A. conceived of the study.

S.M. contributed to electrophysiological recordings.

H.B. contributed to statistical analysis.

D.A.M. performed tissue clearing, lightsheet microscopy, and animal surgeries.

V.A. contributed to MoSeq data processing and study design.

J.N.D. designed the study, performed experiments, analyzed data, and wrote the manuscript.

## Acknowledgments

We thank Kim Loesch, Amy Leonards, Zachary Cantu, Prakruthi Amar Kumar, and Gabrielle Dampf for assistance with animal experiments and care. We thank Dr. Steven Crone for the gift of Chx10-cre mice, Dr. Vicki Tysseling for training on electromyography, Dr. Michael Lane for the gift of PRV-614 and guidance on PRV experiments, Dr. Andrew Murray for providing CVS-N2c rabies and rabies helper virus, Dr. Michael Smotherman for assistance with EMG data analysis, Dr. Richard Gomer for the use of his inverted fluorescence microscope, and Dr. Kumal Sharma for providing the guinea pig Chx10 antibody. We are grateful to the NIDA Drug Supply program for providing us with clozapine-*N*-oxide, and the NIH Virus Center (P40 OD010996) for providing pseudorabies virus. We gratefully acknowledge funding from the Paralyzed Veterans of America Research Foundation (#PVA17_R_0010 to J.N.D.); the Wings for Life Spinal Cord Research Foundation (#237 to A.T.); National Institutes of Health (R01NS116404 to J.N.D., R35GM138098 to H.B., R01NS122961 to D.A.M.); Texas A&M College of Science Undergraduate Research Opportunities Program (to A.B. and K.V.); and Mission Connect, a program of TIRR Foundation (#021-101 to J.N.D.).

