## Supplementary Information for "Functional synaptic connectivity of engrafted spinal cord neurons with locomotor circuitry in the injured spinal cord"

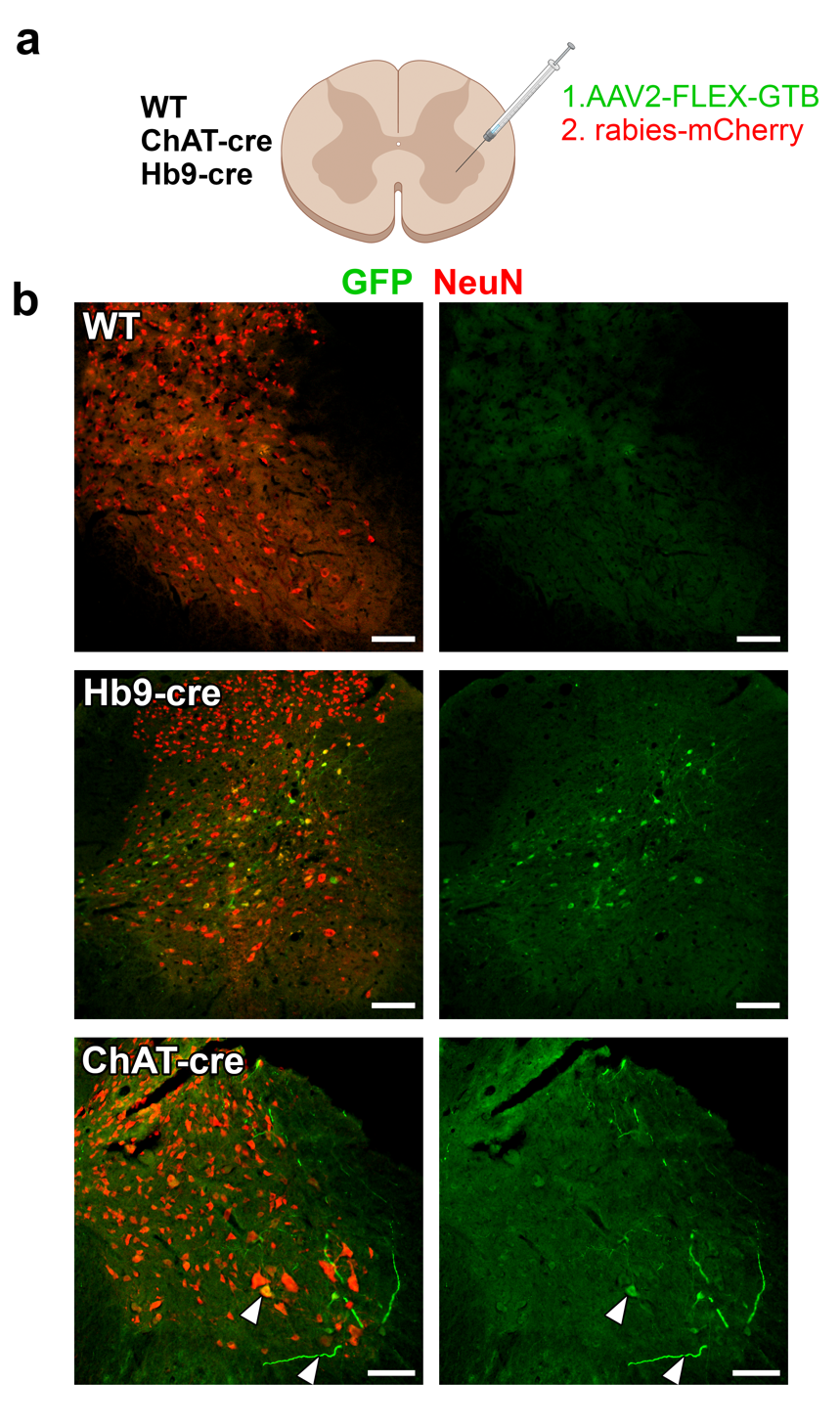

**Supplementary Figure 1. ChAT-cre mice are superior to Hb9-cre mice for targeted transduction of spinal motor neurons.** (**a**) Experimental approach. AAV2-FLEX-GTB (rabies helper + GFP) was injected directly into L2-L5 spinal cord motor pools of either wild-type, ChAT-cre, or Hb9-cre adult mice. Four weeks later, mice were injected with rabies-mCherry. Two weeks later, subjects were sacrificed for histological analysis. (**b**) Transverse images of lumbar (L2-L5) spinal cord four weeks after AAV injection. Rabies-mCherry expression is not shown in this figure in order to highlight Cre-dependent rabies helper expression. No GFP^+^ cells are observed in wild-type subjects. Many GFP^+^ neurons are present in the intermediate gray matter of Hb9-cre subjects, and large-diameter GFP^+^ neurons with motor neuron morphology within the ventral motor pools (arrowheads) are present in ChAT-cre subjects. The ChAT-cre image is the same image used in Figure 1b. Scale bars = 100 μm. Figure was created with Biorender.com.

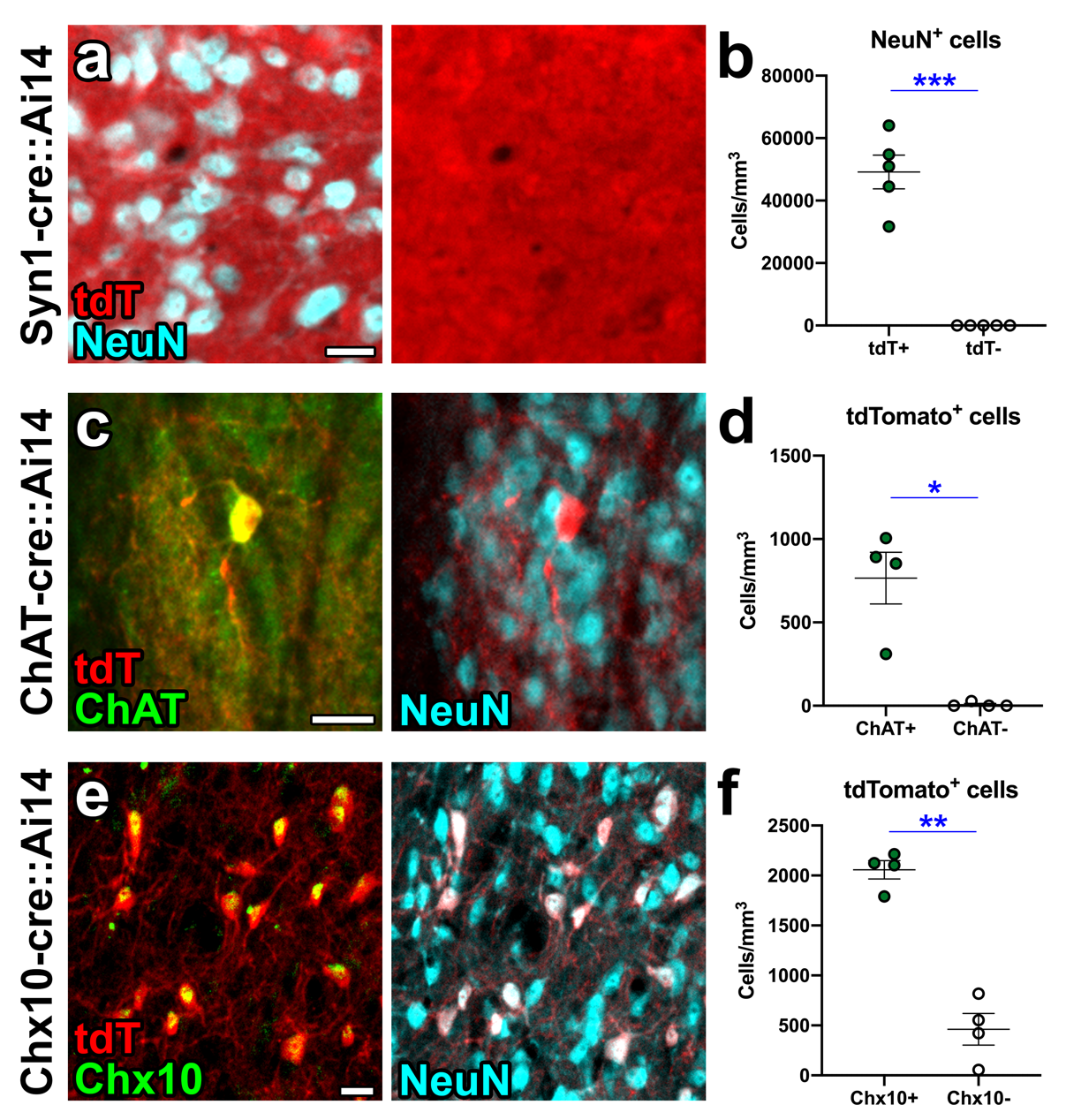

**Supplementary Figure 2. Validation of Cre-dependent transgene expression in neural progenitor cell grafts.** (**a, c, e**) High-magnification images of (**a**) Syn1-cre::Ai14, (**c**) ChAT-cre::Ai14, and (**e**) Chx10-cre::Ai14 NPC grafts at 4 weeks post-transplantation into the host spinal cord. Cre-dependent tdTomato expression is shown in red. (**b**) Quantification of the density of NeuN^+^ neurons in n=4 Syn1-cre::Ai14 grafts that are either tdTomato^+^ or tdTomato^-^. (**d**) Quantification of the density of tdTomato^+^ neurons in n=4 ChAT-cre::Ai14 grafts that are either ChAT^+^ or ChAT^-^. (**f**) Quantification of the density of tdTomato^+^ neurons in n=4 Chx10-cre::Ai14 grafts that are either Chx10^+^ or Chx10^-^ based on immunoreactivity. *p<0.05, **p<0.01, ***p<0.001 by paired, two-tailed t-test. All data are mean ± SEM. Scale bars = 20 μm.

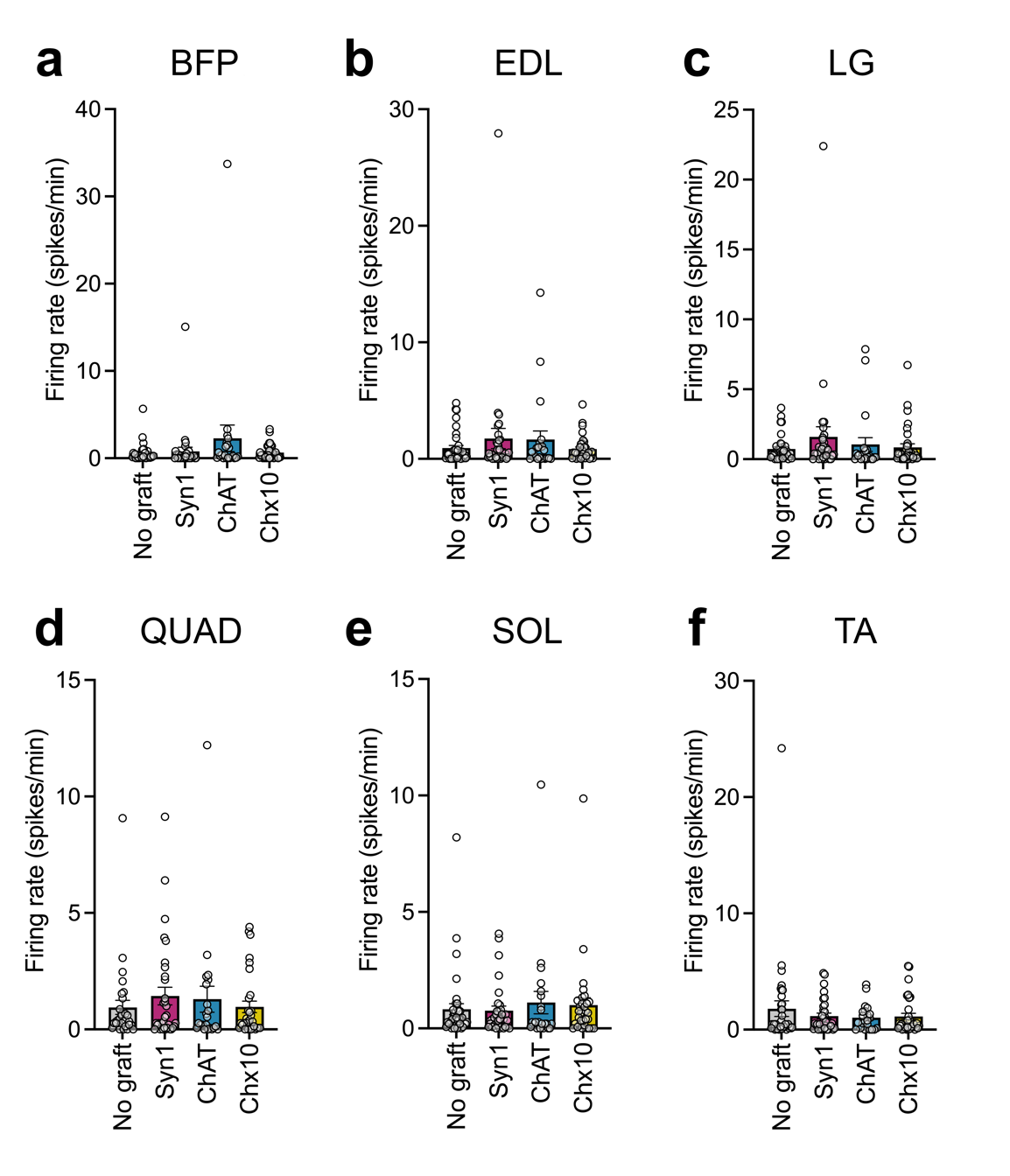

**Supplementary Figure 3. Baseline activity of hindlimb muscles prior to CNO administration.** Muscle activation rate for individual muscle groups (BFP: biceps femoris posterior; EDL: extensor digitorum longus; LG: lateral gastrocnemius; QUAD: quadriceps; SOL: soleus; TA: tibialis anterior) during a 15-minute recording period prior to CNO delivery. Each data point represents an individual subject.

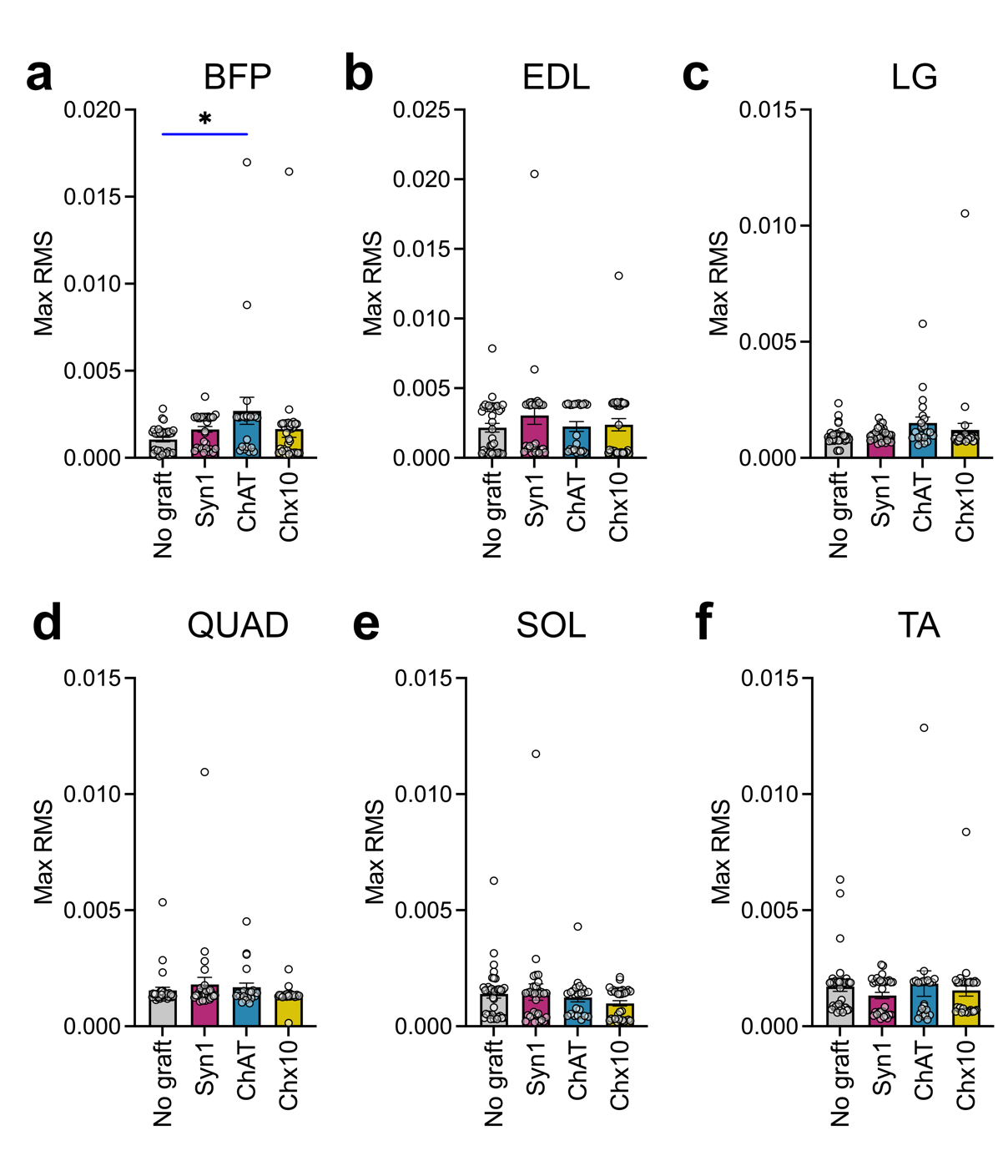

**Supplementary Figure 4. Maximum root mean square for individual muscles post-CNO administration.** Maximum RMS values for individual muscle groups (BFP: biceps femoris posterior; EDL: extensor digitorum longus; LG: lateral gastrocnemius; QUAD: quadriceps; SOL: soleus; TA: tibialis anterior) during the recording period following CNO delivery. Each data point represents an individual subject. *p<0.05 by one way ANOVA + Dunnett’s multiple comparisons test.

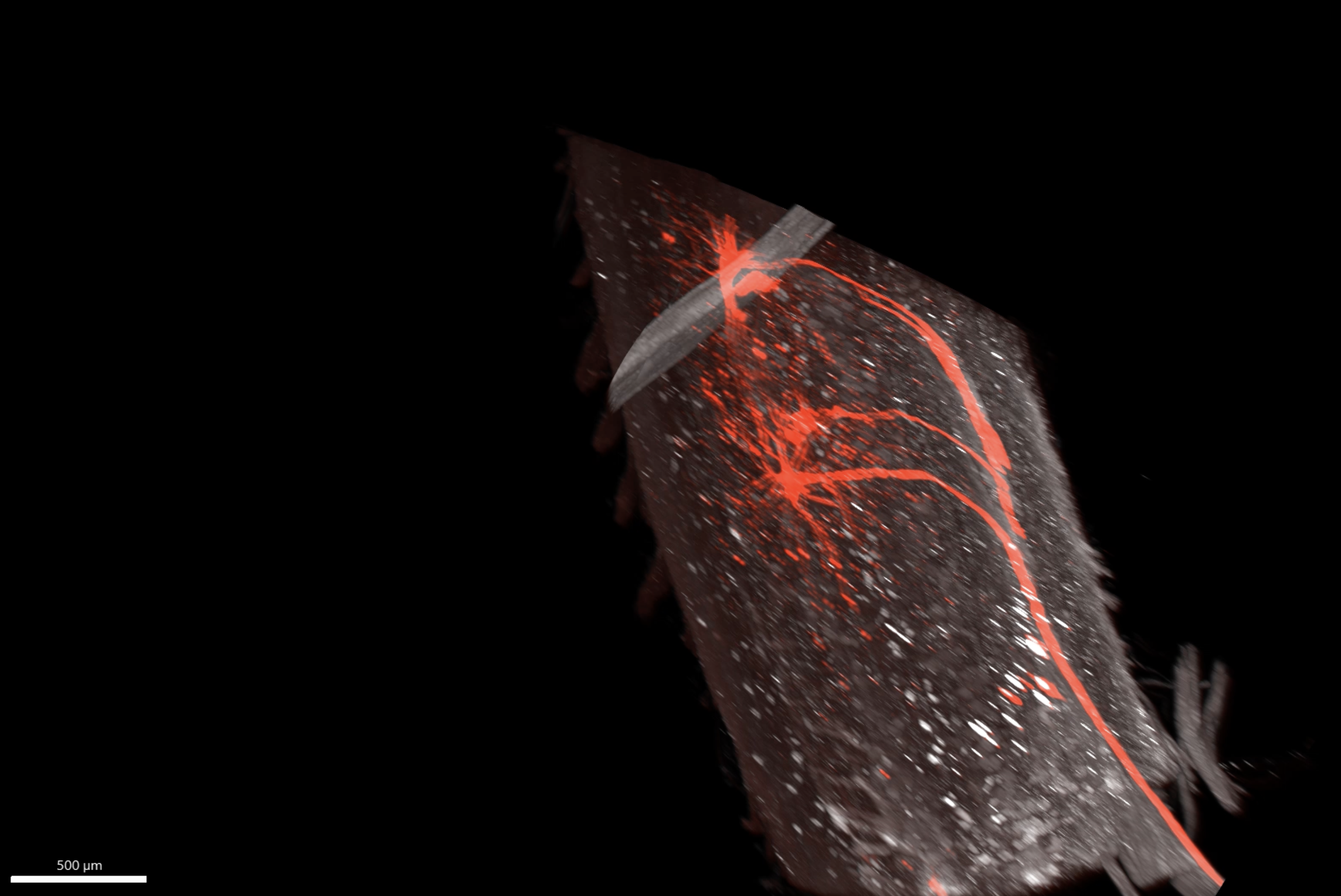

**Supplementary Movie 1.**

Cleared lumbar spinal cord segment taken from an uninjured mouse at 72 hours after PRV-RFP injection into unilateral sciatic nerve. Cleared tissue is immunolabeled with anti-ChAT (white) and anti-RFP (red). Scale bar = 500 μm.

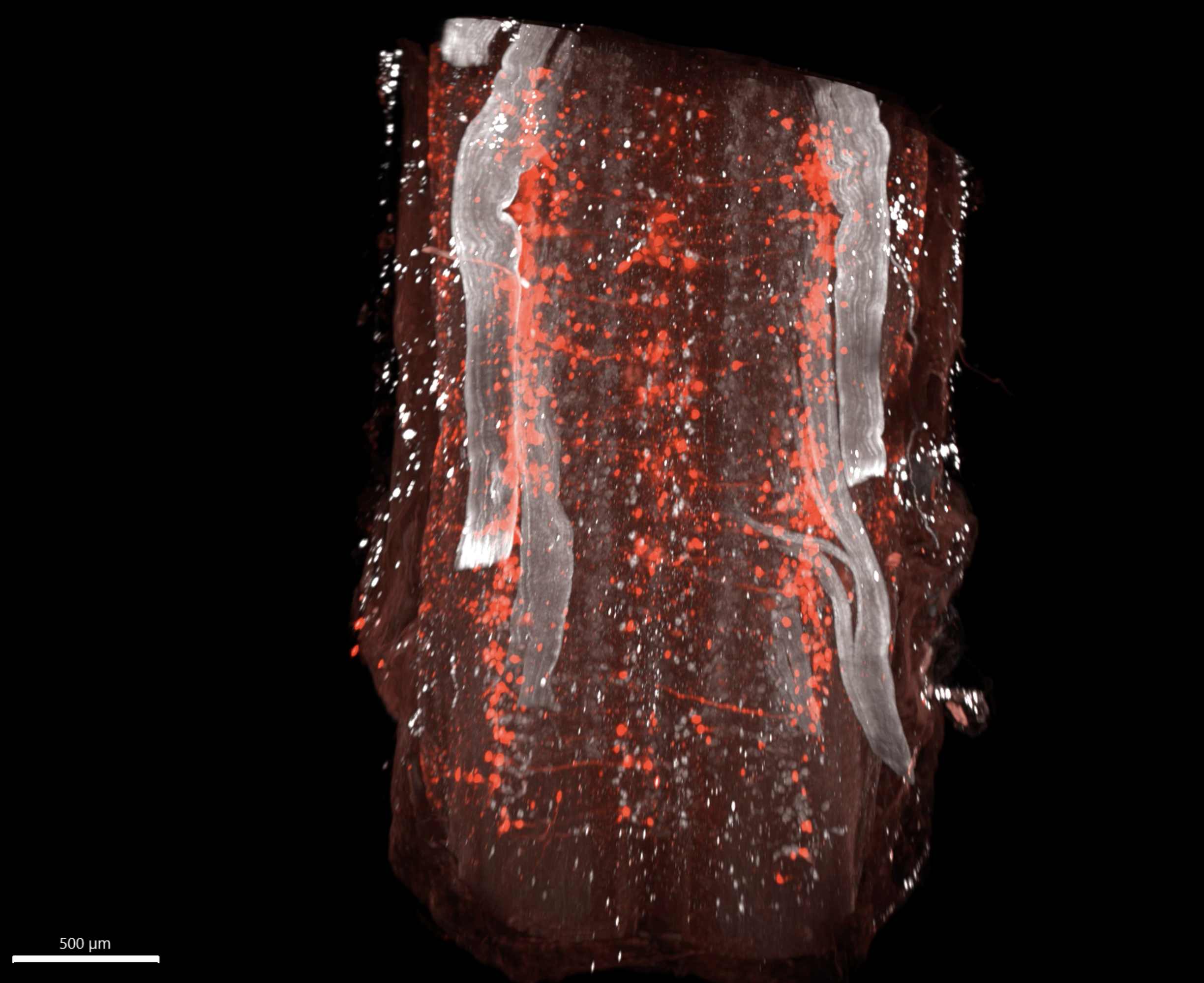

**Supplementary Movie 2.**

Cleared thoracolumbar spinal cord segment taken from an uninjured mouse at 72 hours after PRV-RFP injection into bilateral sciatic nerves. Cleared tissue is immunolabeled with anti-ChAT (white) and anti-RFP (red). Scale bar = 500 μm.

**Supplementary Table 1. Primary and secondary antibodies used in this study.**

| **Antibody** | **Catalog #** | **RRID** | **Dilution** |
| --- | --- | --- | --- |
| **Primary antibodies** | | | |
| Chicken anti-GFP | Aves #GFP-1020 | AB_10000240 | 1:1500 |
| Goat anti-mCherry | Sicgen #AB0040 | AB_2333093 | 1:1500 |
| Rabbit anti-CAMKII | Genetex #GTX135117 | AB_2887447 | 1:250 |
| Rabbit anti-GAD65 + GAD67 | Abcam #ab183999 | AB_2924196 | 1:400 |
| Rabbit anti-Choline acetyltransferase (ChAT) | Genetex #GTX113164 | AB_1949973 | 1:400 |
| Guinea pig anti-Chx10 | Gifted from Kumal Sharma (University of Illinois College of Medicine) |  | 1:200 |
| Guinea pig anti-NeuN | Millipore #ABN90 | AB_11205592 | 1:1000 |
| Rabbit anti-Parvalbumin | Swant #PV27 | AB_2631173 | 1:400 |
| Mouse anti-Calbindin D-28k | Swant #cb300 | AB_3542811 | 1:400 |
| Rabbit anti-cFos | Millipore #ABE457 | AB_2631318 | 1:1000 |
| **Secondary antibodies** |  |  |  |
| Donkey anti-chicken 488 | Jackson ImmunoResearch #703-545-155 | AB_2340375 | 1:1000 |
| Donkey anti-goat Cy3 | Jackson ImmunoResearch #705-165-003 | AB_2340411 | 1:1000 |
| Donkey anti-rabbit 647 | Jackson ImmunoResearch #711-605-152 | AB_2492288 | 1:1000 |
| Donkey anti-guinea pig 647 | Jackson ImmunoResearch #706-605-148 | AB_2340476 | 1:1000 |
| Donkey anti-mouse 647 | Jackson ImmunoResearch #712-605-153 | AB_2340694 | 1:1000 |

**Supplementary Table 2. Detailed description of statistical analyses used in this study.**

| **Fig.** | **Parameter** | **Group size (n)** | **Statistical test** | **Significance level** |
| --- | --- | --- | --- | --- |
| **1i** | Graft volume vs. RABV^+^ neuron density | n=22 | Simple linear regression analysis | R^2^=3.005 x 10^-5^  p=0.9807 |
| **1j** | Graft axon extension at 1500 μm caudal to graft vs. RABV^+^ neuron density | n=21 | Simple linear regression analysis | R^2^=0.2193  p=0.03223 |
| **1m** | Basso Mouse Scale locomotor scores | n=16 (no graft); n=25 (NPC graft) | Two-way repeated measures ANOVA with Sidak’s multiple comparisons test | Time x Treatment main effect: F(13, 507)=0.3014, p=0.9918  Time main effect: F(1.858, 72.47)=1173, p<0.0001  Treatment main effect: F(1, 39)=0.2515, p=0.6188  Subject main effect: F(39, 507)=12.03, p<0.0001  No significant differences between treatment groups |
| **2k** | Graft volume vs. PRV^+^ neuron density | n=19 | Simple linear regression analysis | R^2^=0.2017  p=0.0537 |
| **2l** | Graft axon extension at 1000 μm caudal to graft vs. PRV^+^ neuron density | n=19 | Simple linear regression analysis | R^2^=0.04137  p=0.4036 |
| **3e** | Axon extension from Syn1-cre, ChAT-cre, and Chx10-cre grafts | Syn1-cre (n=11);  ChAT-cre (n=10);  Chx10-cre (n=13) | Two-way repeated measures ANOVA with Tukey’s multiple comparisons test | Distance x Graft type main effect: F(18, 279)=6.610, p<0.0001  Distance main effect: F(1.161, 35.99)=14.97, p=0.0002  Graft type main effect: F(2, 31)=9.866, p=0.0005  Subject main effect: F(31, 279)=18.10, p<0.0001  250 µm:  Syn1-cre vs. ChAT-cre: p=0.0274  Syn1-cre vs. Chx10-cre: p=0.0392  ChAT-cre vs. Chx10-cre: p=0.7052  500 µm:  Syn1-cre vs. ChAT-cre: p=0.0230  Syn1-cre vs. Chx10-cre: p=0.0339  ChAT-cre vs. Chx10-cre: p=0.5686  750 µm:  Syn1-cre vs. ChAT-cre: p=0.0307  Syn1-cre vs. Chx10-cre: p=0.0371  ChAT-cre vs. Chx10-cre: p=0.7819  1000 µm:  Syn1-cre vs. ChAT-cre: p=0.0518  Syn1-cre vs. Chx10-cre: p=0.0732  ChAT-cre vs. Chx10-cre: p=0.6574  1250 µm:  Syn1-cre vs. ChAT-cre: p=0.0454  Syn1-cre vs. Chx10-cre: p=0.0539  ChAT-cre vs. Chx10-cre: p=0.7857  1500 µm:  Syn1-cre vs. ChAT-cre: p=0.0342  Syn1-cre vs. Chx10-cre: p=0.0564  ChAT-cre vs. Chx10-cre: p=0.2968  1750 µm:  Syn1-cre vs. ChAT-cre: p=0.0131  Syn1-cre vs. Chx10-cre: p=0.0146  ChAT-cre vs. Chx10-cre: p=0.9386    2000 µm:  Syn1-cre vs. ChAT-cre: p=0.0498  Syn1-cre vs. Chx10-cre: p=0.0522  ChAT-cre vs. Chx10-cre: p=0.9640  2250 µm:  Syn1-cre vs. ChAT-cre: p=0.0393  Syn1-cre vs. Chx10-cre: p=0.0528  ChAT-cre vs. Chx10-cre: p=0.8274  2500 µm:  Syn1-cre vs. ChAT-cre: p=0.0307  Syn1-cre vs. Chx10-cre: p=0.0307 |
| **3i** | Synaptic punctae density by lamina | Syn1-cre (n=11);  ChAT-cre (n=10);  Chx10-cre (n=13) | Chi-Square followed by Monte Carlo | L3: X-squared=56.134, df=NA, p=0.0004998  L4: X-squared=24.354, df=NA, p=0.04048  L5: X-squared=12.347, df=NA, p=0.6457  L6: X-squared=21.23, df=NA, p=0.09895 |
| **4c** | cFos^+^ cell density in grafts | n=3 (vehicle); n=2 (CNO) | Unpaired, two-tailed t test | t=3.741, df=3  p=0.0333 |
| **4d** | Graft firing rate time series | n=36 neurons (GFP graft); n=47 neurons (Syn1-cre::hM3Dq graft) | Two-way repeated measures ANOVA | Time x Graft type main effect: F(113, 9605)=8.175, p<0.0001  Time main effect: F(3.152, 267.9)=4.995, p=0.0018  Graft type main effect: F(1, 85)=0.3721, p=0.5435  Neuron main effect: F(85, 9605)=166.5, p<0.0001 |
| **4e** | Graft firing rate averages (pre-CNO vs. post-CNO) | n=36 neurons (GFP graft); n=47 neurons (Syn1-cre::hM3Dq graft) | Two-way repeated measures ANOVA | Time x Graft type main effect: F(1, 81)=13.82, p=0.0004  Time main effect: F(1, 81)=11.77, p=0.0009  Graft type main effect: F(1, 81)=0.1158, p=0.7346  Neuron main effect: F(81, 81)=1.617, p=0.0160 |
| **5b** | pre-CNO vs. post-CNO BMS scores | n=18 (no graft); n=14 (Syn1-cre graft); n=12 (ChAT-cre graft); n=17 (Chx10-cre graft) | Paired, two-tailed t test | No graft:  t=0.7656, df=17  p=0.4544  Syn1-cre graft:  t=0.4341, df=13  p=0.6714  ChAT-cre graft:  t=1.000, df=11  p=0.3388  Chx10-cre graft:  t=0.5649, df=15  p=0.5805 |
| **5f** | Maximum firing rate for biceps femoris posterior (BFP) | n=32 (no graft); n=32 (Syn1-cre graft); n=22 (ChAT-cre graft); n=34 (Chx10-cre graft) | One-way ANOVA with Dunnett’s multiple comparisons test | Main effect of treatment: F(3, 116)=1.189, p=0.3170  No graft vs. Syn1-cre graft: p=0.1672  No graft vs. ChAT-cre graft: p=0.8706  No graft vs. Chx10-cre graft: p=0.5299 |
| **5g** | Maximum firing rate for extensor digitorum longus (EDL) | n=36 (no graft); n=32 (Syn1-cre graft); n=22 (ChAT-cre graft); n=34 (Chx10-cre graft) | One-way ANOVA with Dunnett’s multiple comparisons test | Main effect of treatment: F(3, 120)=2.113, p=0.1021  No graft vs. Syn1-cre graft: p=0.1672  No graft vs. ChAT-cre graft: p=0.8706  No graft vs. Chx10-cre graft: p=0.5299 |
| **5h** | Maximum firing rate for lateral gastrocnemius (LG) | n=37 (no graft); n=31 (Syn1-cre graft); n=21 (ChAT-cre graft); n=34 (Chx10-cre graft) | One-way ANOVA with Dunnett’s multiple comparisons test | Main effect of treatment: F(3, 119)=7.673, p<0.0001  No graft vs. Syn1-cre graft: p=0.9584  No graft vs. ChAT-cre graft: p<0.0001  No graft vs. Chx10-cre graft: p=0.8556 |
| **5i** | Maximum firing rate for quadriceps (QUAD) | n=31 (no graft); n=32 (Syn1-cre graft); n=22 (ChAT-cre graft); n=31 (Chx10-cre graft) | One-way ANOVA with Dunnett’s multiple comparisons test | Main effect of treatment: F(3, 112)=1.328, p=0.2690  No graft vs. Syn1-cre graft: p=0.2770  No graft vs. ChAT-cre graft: p=0.3407  No graft vs. Chx10-cre graft: p=0.9898 |
| **5j** | Maximum firing rate for soleus (SOL) | n=38 (no graft); n=32 (Syn1-cre graft); n=22 (ChAT-cre graft); n=32 (Chx10-cre graft) | One-way ANOVA with Dunnett’s multiple comparisons test | Main effect of treatment: F(3, 120)=2.743, p=0.0462  No graft vs. Syn1-cre graft: p=0.0601  No graft vs. ChAT-cre graft: p=0.0400  No graft vs. Chx10-cre graft: p=0.4489 |
| **5k** | Maximum firing rate for tibialis anterior (TA) | n=37 (no graft); n=32 (Syn1-cre graft); n=22 (ChAT-cre graft); n=31 (Chx10-cre graft) | One-way ANOVA with Dunnett’s multiple comparisons test | Main effect of treatment: F(3, 118)=1.596, p=0.1940  No graft vs. Syn1-cre graft: p=0.1237  No graft vs. ChAT-cre graft: p=0.7760  No graft vs. Chx10-cre graft: p=0.2397 |
| **S2b** | tdTomato^+^ vs. tdTomato^-^ neuron density in Syn1-cre::Ai14 grafts | n=5 | Paired, two-tailed t test | t=9.107, df=4  p=0.0008 |
| **S2d** | ChAT^+^ vs ChAT^-^ tdTomato^+^ cell density in ChAT-cre::Ai14 grafts | n=4 | Paired, two-tailed t test | t=5.005, df=3  p=0.0154 |
| **S2f** | Chx10^+^ vs Chx10^-^ tdTomato^+^ cell density in Chx10-cre::Ai14 grafts | n=4 | Paired, two-tailed t test | t=9.445, df=3  p=0.0025 |
| **S3a** | Baseline firing rate for biceps femoris posterior (BFP) | n=32 (no graft); n=32 (Syn1-cre graft); n=22 (ChAT-cre graft); n=34 (Chx10-cre graft) | One-way ANOVA with Dunnett’s multiple comparisons test | Main effect of treatment: F(3, 116)=1.364, p=0.2572  No graft vs. Syn1-cre graft: p=0.9909  No graft vs. ChAT-cre graft: p=0.1798  No graft vs. Chx10-cre graft: p=0.9995 |
| **S3b** | Baseline firing rate for extensor digitorum longus (EDL) | n=36 (no graft); n=32 (Syn1-cre graft); n=22 (ChAT-cre graft); n=34 (Chx10-cre graft) | One-way ANOVA with Dunnett’s multiple comparisons test | Main effect of treatment: F(3, 120)=0.7826, p=0.5059  No graft vs. Syn1-cre graft: p=0.5495  No graft vs. ChAT-cre graft: p=0.7050  No graft vs. Chx10-cre graft: p=0.9988 |
| **S3c** | Baseline firing rate for lateral gastrocnemius (LG) | n=37 (no graft); n=31 (Syn1-cre graft); n=21 (ChAT-cre graft); n=34 (Chx10-cre graft) | One-way ANOVA with Dunnett’s multiple comparisons test | Main effect of treatment: F(3, 119)=0.8262, p=0.4819  No graft vs. Syn1-cre graft: p=0.3345  No graft vs. ChAT-cre graft: p=0.9357  No graft vs. Chx10-cre graft: p=0.9946 |
| **S3d** | Baseline firing rate for quadriceps (QUAD) | n=31 (no graft); n=32 (Syn1-cre graft); n=22 (ChAT-cre graft); n=31 (Chx10-cre graft) | One-way ANOVA with Dunnett’s multiple comparisons test | Main effect of treatment: F(3, 112)=0.4674, p=0.7056  No graft vs. Syn1-cre graft: p=0.6337  No graft vs. ChAT-cre graft: p=0.8653  No graft vs. Chx10-cre graft: p>0.9999 |
| **S3e** | Baseline firing rate for soleus (SOL) | n=38 (no graft); n=32 (Syn1-cre graft); n=22 (ChAT-cre graft); n=32 (Chx10-cre graft) | One-way ANOVA with Dunnett’s multiple comparisons test | Main effect of treatment: F(3, 120)=0.2576, p=0.8558  No graft vs. Syn1-cre graft: p=0.9977  No graft vs. ChAT-cre graft: p=0.8638  No graft vs. Chx10-cre graft: p=0.9505 |
| **S3f** | Baseline firing rate for tibialis anterior (TA) | n=37 (no graft); n=32 (Syn1-cre graft); n=22 (ChAT-cre graft); n=32 (Chx10-cre graft) | One-way ANOVA with Dunnett’s multiple comparisons test | Main effect of treatment: F(3, 119)=0.6421, p=0.5894  No graft vs. Syn1-cre graft: p=0.6114  No graft vs. ChAT-cre graft: p=0.5465  No graft vs. Chx10-cre graft: p=0.5759 |
| **S4a** | Maximum root mean square for biceps femoris posterior (BFP) | n=32 (no graft); n=32 (Syn1-cre graft); n=22 (ChAT-cre graft); n=34 (Chx10-cre graft) | One-way ANOVA with Dunnett’s multiple comparisons test | Main effect of treatment: F(3, 116)=2.427, p=0.0691  No graft vs. Syn1-cre graft: p=0.5928  No graft vs. ChAT-cre graft: p=0.0226  No graft vs. Chx10-cre graft: p=0.5637 |
| **S4b** | Maximum root mean square for extensor digitorum longus (EDL) | n=35 (no graft); n=32 (Syn1-cre graft); n=22 (ChAT-cre graft); n=34 (Chx10-cre graft) | One-way ANOVA with Dunnett’s multiple comparisons test | Main effect of treatment: F(3, 119)=0.7572, p=0.5203  No graft vs. Syn1-cre graft: p=0.3824  No graft vs. ChAT-cre graft: p=0.9992  No graft vs. Chx10-cre graft: p=0.9779 |
| **S4c** | Maximum root mean square for lateral gastrocnemius (LG) | n=37 (no graft); n=31 (Syn1-cre graft); n=21 (ChAT-cre graft); n=34 (Chx10-cre graft) | One-way ANOVA with Dunnett’s multiple comparisons test | Main effect of treatment: F(3, 119)=1.545, p=0.2064  No graft vs. Syn1-cre graft: p=0.9923  No graft vs. ChAT-cre graft: p=0.1253  No graft vs. Chx10-cre graft: p=0.6051 |
| **S4d** | Maximum root mean square for quadriceps (QUAD) | n=31 (no graft); n=32 (Syn1-cre graft); n=22 (ChAT-cre graft); n=31 (Chx10-cre graft) | One-way ANOVA with Dunnett’s multiple comparisons test | Main effect of treatment: F(3, 112)=1.069, p=0.3655  No graft vs. Syn1-cre graft: p=0.6576  No graft vs. ChAT-cre graft: p=0.9414  No graft vs. Chx10-cre graft: p>0.7957 |
| **S4e** | Maximum root mean square for soleus (SOL) | n=38 (no graft); n=32 (Syn1-cre graft); n=22 (ChAT-cre graft); n=32 (Chx10-cre graft) | One-way ANOVA with Dunnett’s multiple comparisons test | Main effect of treatment: F(3, 120)=0.9038, p=0.4415  No graft vs. Syn1-cre graft: p=0.9884  No graft vs. ChAT-cre graft: p=0.9487  No graft vs. Chx10-cre graft: p=0.4289 |
| **S4f** | Maximum root mean square for tibialis anterior (TA) | n=37 (no graft); n=32 (Syn1-cre graft); n=22 (ChAT-cre graft); n=32 (Chx10-cre graft) | One-way ANOVA with Dunnett’s multiple comparisons test | Main effect of treatment: F(3, 119)=0.6133, p=0.6077  No graft vs. Syn1-cre graft: p=0.5998  No graft vs. ChAT-cre graft: p=0.9811  No graft vs. Chx10-cre graft: p=0.9500 |
